# Cell Simulation as Cell Segmentation

**DOI:** 10.1101/2024.04.25.591218

**Authors:** Daniel C. Jones, Anna E. Elz, Azadeh Hadadianpour, Heeju Ryu, David R. Glass, Evan W. Newell

## Abstract

Single-cell spatial transcriptomics promises a highly detailed view of a cell’s transcriptional state and microenvironment, yet inaccurate cell segmentation can render this data murky by misattributing large numbers of transcripts to nearby cells or conjuring nonexistent cells. We adopt methods from ab initio cell simulation to rapidly infer morphologically plausible cell boundaries that preserve cell type heterogeneity. Benchmarking applied to datasets generated by three commercial platforms show superior performance and computational efficiency of this approach compared with existing methods. We show that improved accuracy in cell segmentation aids greatly in detection of difficult to accurately segment tumor infiltrating immune cells such as neutrophils and T cells. Lastly, through improvements in our ability to delineate subsets of tumor infiltrating T cells, we show that CXCL13-expressing CD8+ T cells tend to be more closely associated with tumor cells than their CXCL13-negative counterparts in data generated from renal cell carcinoma patient samples.

## 1 Introduction

Recent imaging-based in-situ spatial transcriptomics technologies [Chen et al., 2015, He et al., 2022, Janesick et al., 2022] offer a highly detailed readout of the locations of RNA transcripts from a gene panel of several hundred, and in some cases, thousands of genes. To take full advantage of this data, transcripts must first be attributed to cells by segmenting space into regions occupied by each cell. Cell segmentation is in some ways analogous to read alignment scRNA-Seq; a computationally intensive pre-processing task that is necessary to arrive at any measurement of per-cell gene expression. Yet unlike read alignment, transcripts are misassigned at an alarming rate due to inaccuracies in segmentation dramatically shaping the interpretation of the data.

If mis-segmentation merely injected some amount of uncorrelated noise into the data, it would be regrettable, but no real cause to question results from downstream analysis. What makes it pernicious, and a cause for deep concern is that by misattributing transcripts from adjacent cells, cell proximity and gene expression become confounded in the data in a way that most statistical analyses are entirely unequipped to account for. Since studying the relationship between gene expression and spatial organization is arguably the entire point of spatial transcriptomics, there could hardly be a more unfortunate type of bias to afflict the data with than to confound the two.

It calls into question differential expression calls, as a gene can appear upregulated in some cell population simply by being nearer to another cell population expressing that gene. Differential gene expression tests for count data, mostly developed for RNA-Seq, can account for noise in the form of drop-out or overdispersion, but have no concept of noise from mis-segmentation. The result is numerous false positives. For example, lymphocytes recruited to a tumor region could appear to be expressing an unusual collection of genes, but to determine which of these expression changes represent potentially interesting biology, we first need to rule out those whose transcripts were erroneously misattributed from the adjacent tumor.

Even labeling cell types is burdened with systemic error and fraught with ambiguity. The obvious result of mixing signals of adjacent cells is the introduction of large numbers of cells of artificially hybrid type that may be impossible to identify with any confidence or may naively appear to be novel cell types. Less widely appreciated is the degree to which this signal mixing introduces asymmetrical bias into the data. Because the number of transcripts identified in a cell can differ by one or two orders of magnitude, spillover between a cell with 1000 transcripts and one with 50 transcripts can result in the former’s signal completely dominating the latter’s. If we are lucky this might produce an ambiguous cell, but often the identity of the low population cell is completely buried.

The traditional method of cell segmentation, and the one deployed out the box by the major in-situ spatial transcriptomics platforms is image-based segmentation. By this approach, a deep learning model is applied to microscopy images with one or more fluorescent stains applied. At minimum, this includes a nuclear stain but often also includes various membrane stains or a nonspecific mRNA stain. With cell boundaries established, transcripts are then trivially assigned to cells by testing which region they fall into.

Many years of development have gone into refining image based cell segmentation, which long predates in-situ spatial transcriptomics. Early success was found using heuristics like the watershed algorithm [Beucher and Meyer, 1992] that operate on single-channel images. More recently, the task has fallen to deep learning models like U-Net [Ronneberger et al., 2015], Mesmer [Greenwald et al., 2021], and Cellpose [Stringer et al., 2021]. As in all deep learning image recognition tasks, the model is only successful to the extent that it was trained on representative data. In the cell segmentation task, this can be problematic, both because cells come in all shapes and sizes, and because different platforms and experiments may use differing staining strategies. Lacking a pre-trained model, a practitioner is forced to either manually label data to train their own, or risk mis-segmentation with a potentially ill-fitting model. Though recent work [Israel et al., 2023] has focused on trying to improve how well models generalize, this remains a major hurdle. The initial release of 10X Genomics Xenium side steps poor generalization by only segmenting nuclei, which are far easier to consistently segment, and then expanding the nuclei by some fixed radius, combining relatively accurate assignment of nuclear transcripts with highly approximate assignment of cytoplasmic transcripts.

Image based cell segmentation must contend with the unfortunate reality that while images are 2D, cells exist in 3D space, and even a 5μm thick section will often have cells that overlap significantly on the z-axis. Regardless of how successful a staining strategy is and sophisticated the deep learning model, we cannot hope to discern a complex 3D boundary between cells with only an image. Multiple images at varying focal lengths provide some amount of 3D information, which can be exploited by interpolating multiple 2D cell boundaries. Cellpose Stringer et al. [2021] proposed an alternative, jointly modeling multiple focal lengths to infer 3D volume, at the cost of dramatically higher computational costs.

Recently more specialized segmentation algorithms have been proposed that seek to exploit the transcript data itself to perform segmentation, rather than relying on corresponding images. Most of these are supervised deep learning models, that otherwise approach the problem from very different perspectives. SCS [Chen et al., 2023] discretizes the transcript data into what resembled image layers, then uses a transformer model to predict a gradient pointing from the cell center. GeneSegNet [Wang et al., 2023] similarly discretizes the data to resemble image layers, stacks these with other image data, and uses a U-Net based model to infer cell masks. BIDCell [Fu et al., 2024] also uses a U-Net based model, but with a different objective function designed to incorporate prior information about cell type and gene expression patterns. Instead of collapsing transcript data into image layers, Bering [Jin et al., 2023] considers the k-nearest neighbor (KNN) graph induces over transcripts. It poses segmentation as a binary classification problem, inferring edges between nearest neighbors to be either intracellular or intercellular using an architecture that combines image data with the KNN graph using a graph convolutional network.

An interesting alternative is Baysor [Petukhov et al., 2022], which adopts a statistical model of gene expression and the spatial distribution of transcripts, treating the problem as a large mixture model and inferring the assignments of transcripts to cells using expectation maximization. This approach is unusual not only because it is not based on deep learning, but because it is entirely unsupervised. This is an appealing aspect which few methods share. SCS attempts to avoid the problem of explicitly labeled training by assuming sample itself contains a large population of unassigned transcripts it can train on as negative examples. BIDCell does not necessitate labeled training data, but does require detailed prior information about gene co-expression patterns to inform its segmentation. As the technology becomes more widespread, it becomes increasingly unrealistic that each experiment will be run with its own bespoke segmentation model, fine-tuned on manually labeled cells.

Here we propose a segmentation method, Proseg (from *probabilistic segmentation*), that is based on an entirely unsupervised probabilistic model of the spatial distribution of transcripts, capable of efficiently and accurately segmenting cells without the need of multimodal staining strategies. The method takes as its inspiration Cellular Potts models (CPMs) [Graner and Glazier, 1992], which simulate rudimentary cellular behaviors on a lattice of pixels or voxels by optimizing a contrived energy function. With Proseg we employ a similar strategy to generate cell morphologies, but in place of a contrived function we use a probabilistic model of the spatial distribution of observed transcripts. Cell morphologies are initialized using a nuclear stain, then expanded and altered at random until they best explain the observed data. Simultaneously, and uniquely to this method, we are also able to reposition transcripts that appear in an implausible position, whether due to RNA leaking from adjacent cells or measurement error.

Deploying this method across three in situ spatial transcriptomics platforms (Vizgen MERSCOPE, NanoString CosMx, and 10X Xenium), we found that Proseg efficiently infers accurate cell segmentation, which can have dramatic effects on the interpretation of the data, dramatically shifting the composition of inferred cell types and patterns of gene coexpression.

## 2 Results

### 2.1 Proseg Model and Implementation

As a starting point we draw on the field of ab initio cell simulation. The Cellular Potts model (CPM) [Graner and Glazier, 1992] represents cells as occupying voxels in a lattice and simulated their behavior by optimizing a contrived objective function, which is designed to induce interesting behaviors analogous to those observed in real cell populations. For example, this function might assign a penalty to contact between cells of different types. The simulation itself then proceeds using a Monte Carlo sampling procedure that repeatedly modifies cell boundaries to minimize this objective function, gradually inducing cell sorting over many iterations (Figure 1a).

**Figure 1:**
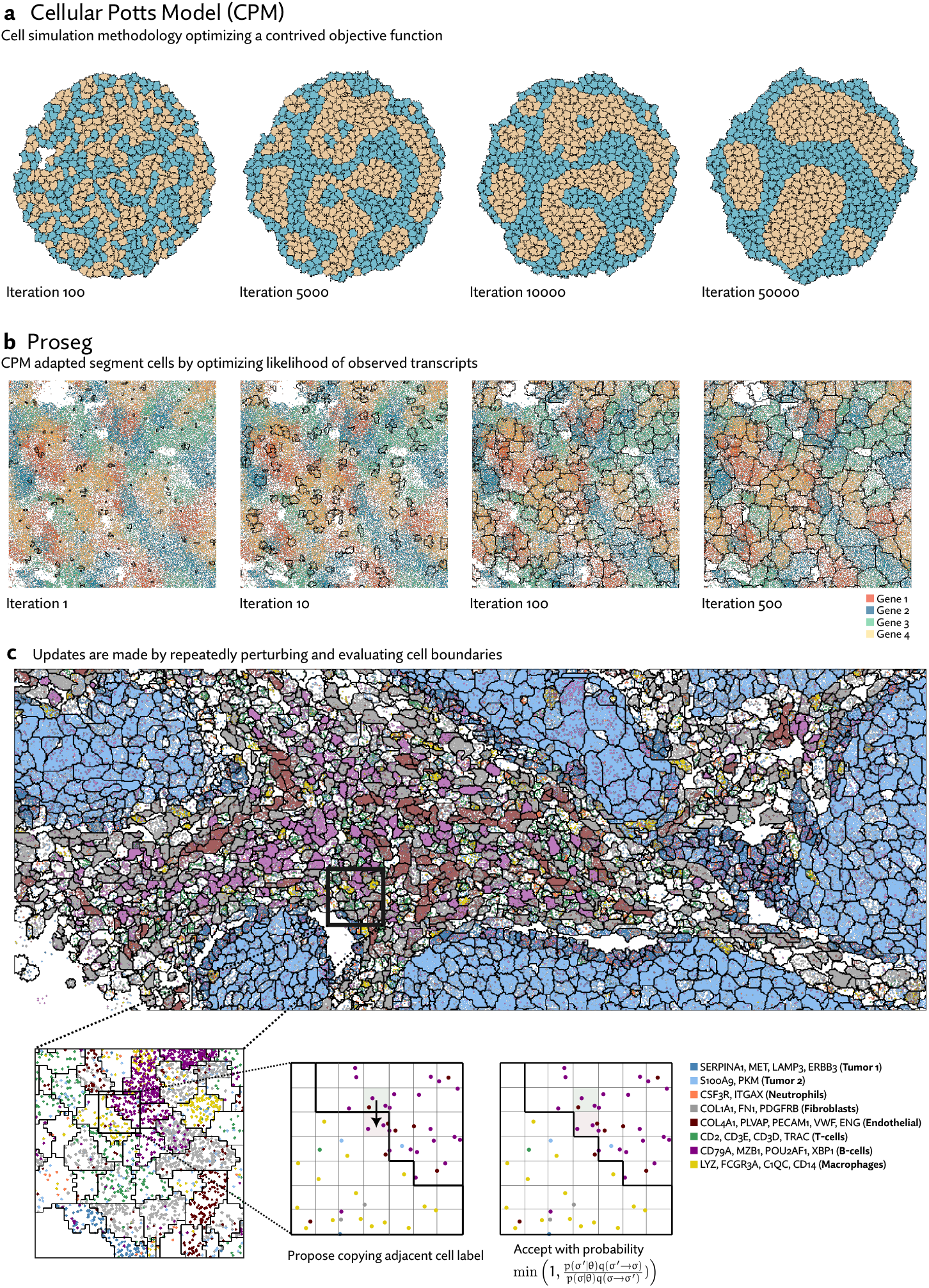
**(a)** Cellular Potts models (CPMs) represent cells on a grid of pixels or voxels and repeatedly perturbs cell boundaries to optimize a contrived objecive function. This function can be designed to induce specific behaviors. In the example shown here, cells have higher affinity to those of the same time, and after many iterations of the simulation, migrate and sort themselves (images generated with Morpheus [Starruß et al., 2014]). **(b)** Proseg (“probabilistic segmentation”) adapts this simulation framework to instead generate cell boundaries that best explain the observed spatial distribution of transcripts, turning a cell simulation methodology into a cell segmentation methodology. In place of a designed objective function a probabilistic model of gene expression is used. In this example, cell boundaries are gradually optimized to explain the distribution of four highly cell type specific genes. **(c)** Proseg and CPMs operate under the same basic sampling framework, demonstrated here in a section of the MERSCOPE dataset. Cell boundaries are perturbed by copying the label of an adjacent voxel. The change in the objective function is evaluated determining whether the perturbation is accepted or rejected. This basic sampling procedure is iterated until convergence.

CPMs have been shown to be highly flexible. For example, in addition to the cell sorting example, they have been used to simulate migration and chemotaxis with an actin-inspired mechanism [Niculescu et al., 2015] and mimic aspects of wound healing [Scianna, 2015]. Furthermore, extremely efficient algorithms exist to run these simulations in a highly parallelized fashion [Chen et al., 2007, Tapia and D’Souza, 2011, Sultan et al., 2023].

Because the simulation is already run using a sampling procedure resembling the Metropolis algorithm, it is a short conceptual leap to convert CPMs into a full-fledged probabilistic model with Markov-chain Monte Carlo based inference. In place of an energy function designed to elicit specific behaviors we use a probabilistic model of gene expression. In short, rather than simulating cells to best satisfy a contrived objective function, Proseg simulates cells to best explain the distribution of observed transcripts (Figure 1b). Despite this change in objective, the same basic CPM sampling procedure is used: random proposals are made by copying the label of one voxel to an adjacent position. This change is evaluated and either accepted or rejected, here according to the Metropolis algorithm (Figure 1c).

Though this avoids the issue of supervision and mismatched models, it creates the risk that elaborate cell morphologies will be generated to engineer cells with artificially parsimonious gene expression — a kind of cellular gerrymandering. We adopt recent advancements in modeling and inference for count data [Zhou and Carin, 2015, Dadaneh et al., 2020] to construct a model that accommodates observed variation in gene expression without coercing cells into predefined types. Combined with techniques from the CPM literature for constraining cellular surface area [Magno et al., 2015], Proseg is informed by gene expression, avoiding egregious misassignment errors, while generating plausible cell morphologies.

Our model can also account for some probability that a transcript is misplaced in a strategy we call *transcript repositioning*. The Proseg sampler, in addition to sampling model parameters and cell boundaries, can propose shifts of transcripts to nearby positions. Mis-placed transcripts can be due to RNA leaking from cells at some rate, a side effect of permeabilization steps most protocols take to grant probes access to cellular mRNA. This phenomenon is clearly observable in the data (Supplementary Figure 12). The scheme can also be justified if we accept that there is some error in reported transcript positions (particularly on the z-axis) and as a means of compensating for the inherent imprecision in cell boundaries that are confined to a lattice of voxels.

Further details of the probabilistic model and implementation are described in the Methods section.

### 2.2 Benchmarking segmentation

To compare segmentation methods, we collected four datasets across three platforms, including Vizgen MERSCOPE, NanoString CosMx, and 10X Xenium with and without their recently released multimodal segmentation method. Three of these datasets are publically available, generated from human lung cancer samples. We additionally generated Xenium data across several renal cell carcinoma (RCC) samples.

Each of these platforms provides by default an image-based segmentation approach applied to multi-channel images, each adopting somewhat different staining strategies. In MERSCOPE and CosMx these are Cellpose models, while 10X uses a custom deep learning model. In the standard version of Xenium, which used to generate the RCC data, only a nuclear stain was used, and cell boundaries were defined heuristically by expanding the inferred nuclear boundaries by a fixed distance. More recently, Xenium offers optionally a “multimodal” segmentation scheme that uses additional stains to better define boundaries.

Lacking a high confidence ground truth segmentation to compare to, we sought to quantify suspicious patterns of co-expression across segmentation methods. We compared rates of co-expression between all pairs of genes in the nuclear segmentation to nuclear segmentation with fixed radius expansion. We selected pairs of genes that showed an over 50% increase in co-expression after nuclei expansion. Though some of these pairs may be explained by subcellular localization to the cytoplasm, the far more common cause is incorrect assignment of nearby transcripts. We observed a consistent reduction in co-expression of these suspicious gene pairs using Proseg, compared to Baysor, Bering, or image-based deep learning methods (Figure 2a). Though not perfect proxy for segmentation accuracy, it does capture the most pernicious form of missegmentation error, namely transcript misassignment, that we would hope to reduce.

**Figure 2:**
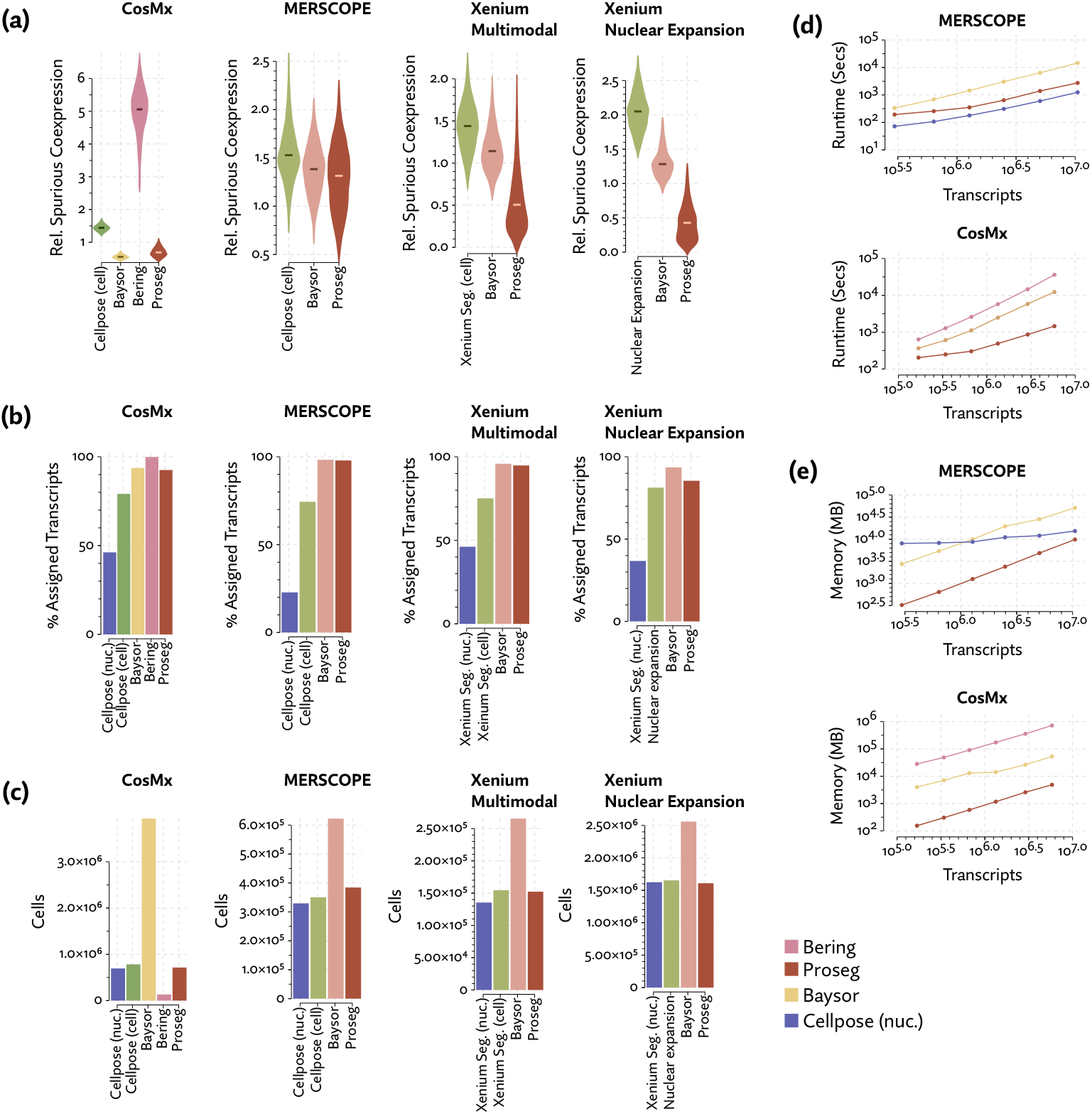
Benchmarking of competing segmentation methods across four spatial transcriptomics datasets. (a) Spuriously co-expressed gene pairs were defined as those with rates of co-expression that increase dramatically when nuclear boundaries are expanded. Relative spurious co-expression rates were computed as the rate of co-expression of these spurious pairs relative to nuclear segmentation, with lower rates suggestive of higher quality segmentation. (b) Image-based segmentation methods fail to assign large portions of the transcripts compared to transcript-driven methods like Bering, Baysor, and Proseg. (c) Compared to Proseg and image-based segmentation, Baysor predicts dramatically more cells, suggesting systematic over-segmentation, while Bering predicted far fewer cells. Comparisons of memory and runtime on (d) MERSCOPE and (e) CosMx datasets show that Proseg is generally an order of magnitude more efficient that Baysor and competitive with Cellpose.

Frequently, image-based segmentation approaches failed to assign a large proportion of the observed transcripts (Figure 2b), suggesting overly conservative cell borders, despite their higher levels of spurious co-expression. These unassigned transcripts are unlikely to be noise. Each platform has negative controls in some capacity, which do not corroborate such a high level of spuriously observed transcripts. Thus, the 20% or more of transcripts left unassigned by each platform’s default method represents a genuine large-scale short-coming in existing image-based segmentation approaches.

Like Proseg, Baysor also consistently improves on the rate spurious co-expression, but it does so at the cost of introducing many more cells (Figure 2c). The Baysor manuscript describes that it will “in some cases, nearly double the number of cells” [Petukhov et al., 2022], which is consistent with our observations, even after excluding large numbers of cells with fewer than 10 assigned transcripts. Yet, the evidence that these cells are real is lacking. Baysor introduces these extra cells to induce parsimonious gene expression distributions, but accepting this we are forced to believe that nearly half of all cells either do not show up on nuclear stains or are missed by other methods. No prior literature suggests that DAPI might fail to stain such a high proportion of cells, and manual inspection of the nuclear segmentation does not lend credence to anything approaching a 40-50% error rate. Meanwhile, Proseg is able to produce similarly well-defined cell types and assign similar numbers of transcripts without the dubious increase in cell count.

Comparing runtime and memory usage (Figure 2de), we found that while all methods scaled roughly linearly with the number of transcripts, Proseg was approximately an order of magnitude faster than Baysor and Bering, though moderately slower Cellpose. Proseg also requires roughly an order of magnitude less memory than Baysor or Bering. Cellpose, which is operating on image tiles rather than transcript tables, has a shallower curve and, for sufficiently large datasets, will require less memory.

### 2.3 Improved resolution of neutrophils and T-cells in MERSCOPE and CosMx lung cancer tumor samples

The MERSCOPE data we evaluated was taken from the Vizgen MERFISH FFPE Immuno-oncology data release [Vizgen, 2022]. The platform provides cell segmentation based on custom trained Cellpose model applied to a cell boundary stain, as well as DAPI and Poly-T stains. We compared this to Cellpose applied just to the DAPI stain. Nuclei are far easier to segment than full cells, so this provides us with incomplete but relatively high confidence transcript assignments. We also ran Baysor, initialized using the Cellpose segmentation, and Proseg initialized by the Cellpose nuclear segmentation. For each method, we discarded cells with fewer than 10 assigned transcripts.

The same major cell type clusters were identifiable under each segmentation method (Figure 3a), to varying degrees of ambiguity. Contributing to this ambiguity, the provided Cellpose model failed to assign roughly 26% of the transcripts, which suggests either very high levels of noise or overly conservative segmentation. Both Baysor and Proseg are able to remedy this, assigning nearly all the transcripts, but differ most significantly in that Baysor introduces an extra 230 thousand cells, a 62% increase over Proseg (Figures 2c, 3b).

**Figure 3:**
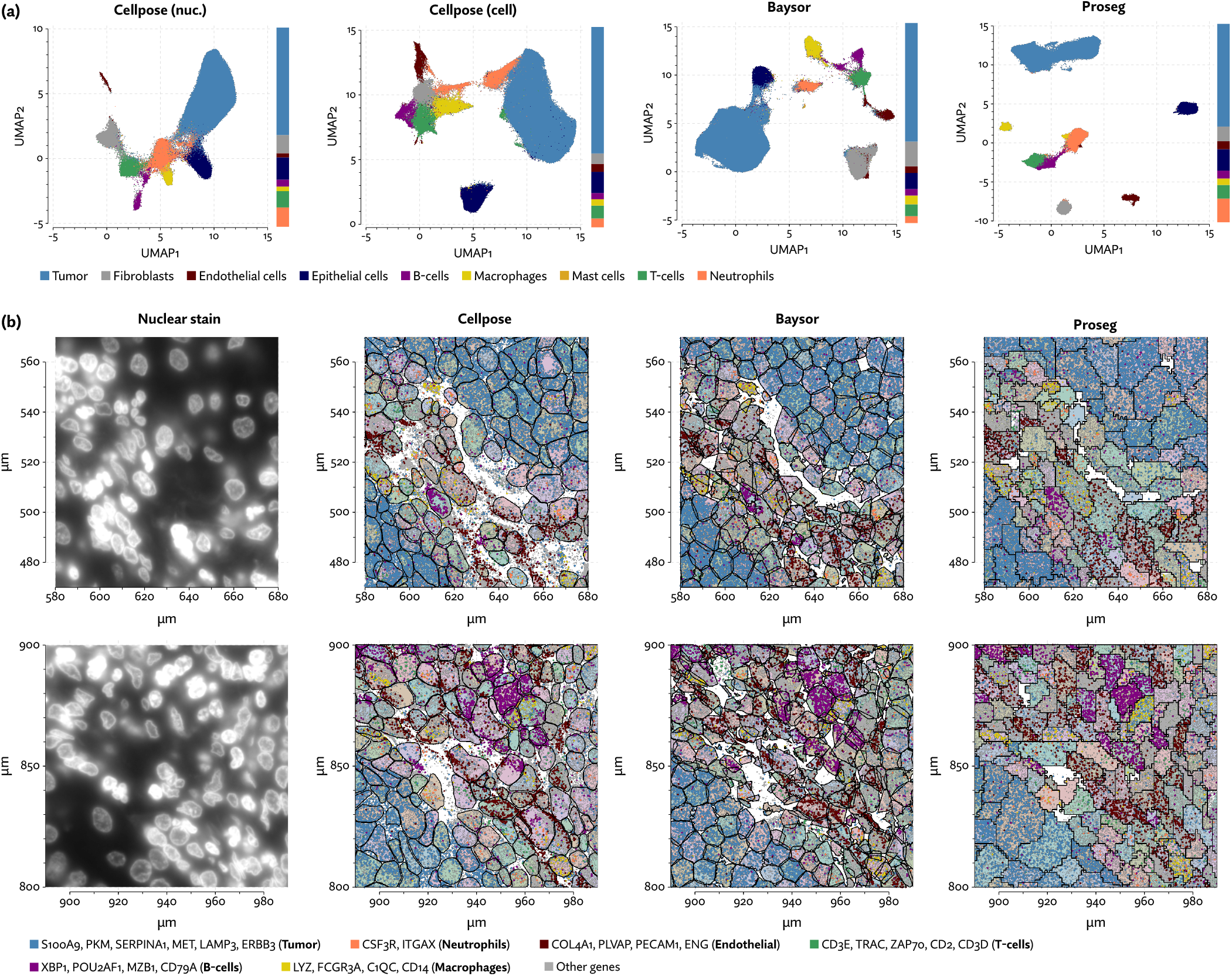
Segmentation results on the MERSCOPE lung cancer dataset. (a) UMAP plots corresponding to each segmentation method. Cell type proportions are shown stacked bar plots on the margins. (b) A comparison of cell segmentation in two regions across three segmentation methods, shown along with specific subsets of highly cell type specific transcripts.

Compared to nuclear segmentation, Cellpose and Baysor introduced significant distortions into the cell type composition (Figure 3a). Most notably, the proportion of neutrophils and T-cells was diminished by nearly half. This is easily explained by the asymmetrical nature of cell segmentation error. T-cells and neutrophils had a median of 81 and 96 assigned transcripts, respectively, in the provided segmentation. Tumor cells had a median of 291 in comparison. When segmentation errors are made between tumor cells and infiltrating neutrophils, the more likely outcome is a neutrophil mislabeled as tumor cell, rather than vice versa, exactly as observed here. Because of this, segmentation by both Cellpose and Baysor dramatically under-represents both infiltrating neutrophils, which are systematically misidentified as tumor cells, and T-cells in the tumor periphery, which are often misidentified as fibroblasts or endothelial cells (Supplementary Figure 1).

Neutrophils were found to be the dominant immune cell type in non-small cell lung cancer and present in high proportion in other lung center types [Kargl et al., 2017]. A later study showed a similarly high proportion of neutrophils in lung squamous cell carcinoma and an overall association between neutrophil content and tumor gene expression heterogeneity [Wu et al., 2021]. Agreeing with this study, we also see high expression of CXCL1 in tumor cells, and high expression of the corresponding receptor CXCR1 in neutrophils, suggesting a mechanism by which neutrophils are recruited into the interior of the tumor. Neutrophil content has been proposed as a variable that is highly predictive of the success or failure of immune checkpoint therapy [Kargl et al., 2019]. Dramatically miscounting neutrophils and T-cells is therefore far from inconsequential.

We next examined a public demonstration dataset provided by NanoString for the CosMx platform [He et al., 2022]. The dataset consists of 5 human lung cancer samples, one of which was run with three replicates, and another with two, for a total of 8 samples (a portion of one sample is shown in Figure 4b). The image-based cell segmentation provided by CosMx is based on a Cellpose model applied to a multichannel image from four fluorescent stains. A small subset of this dataset was examined by a paper describing the Bering segmentation method [Jin et al., 2023]. Though Bering is a supervised model, we can make a fair comparison here by using training data published by the authors to run segmentation on the entirety of the dataset.

**Figure 4:**
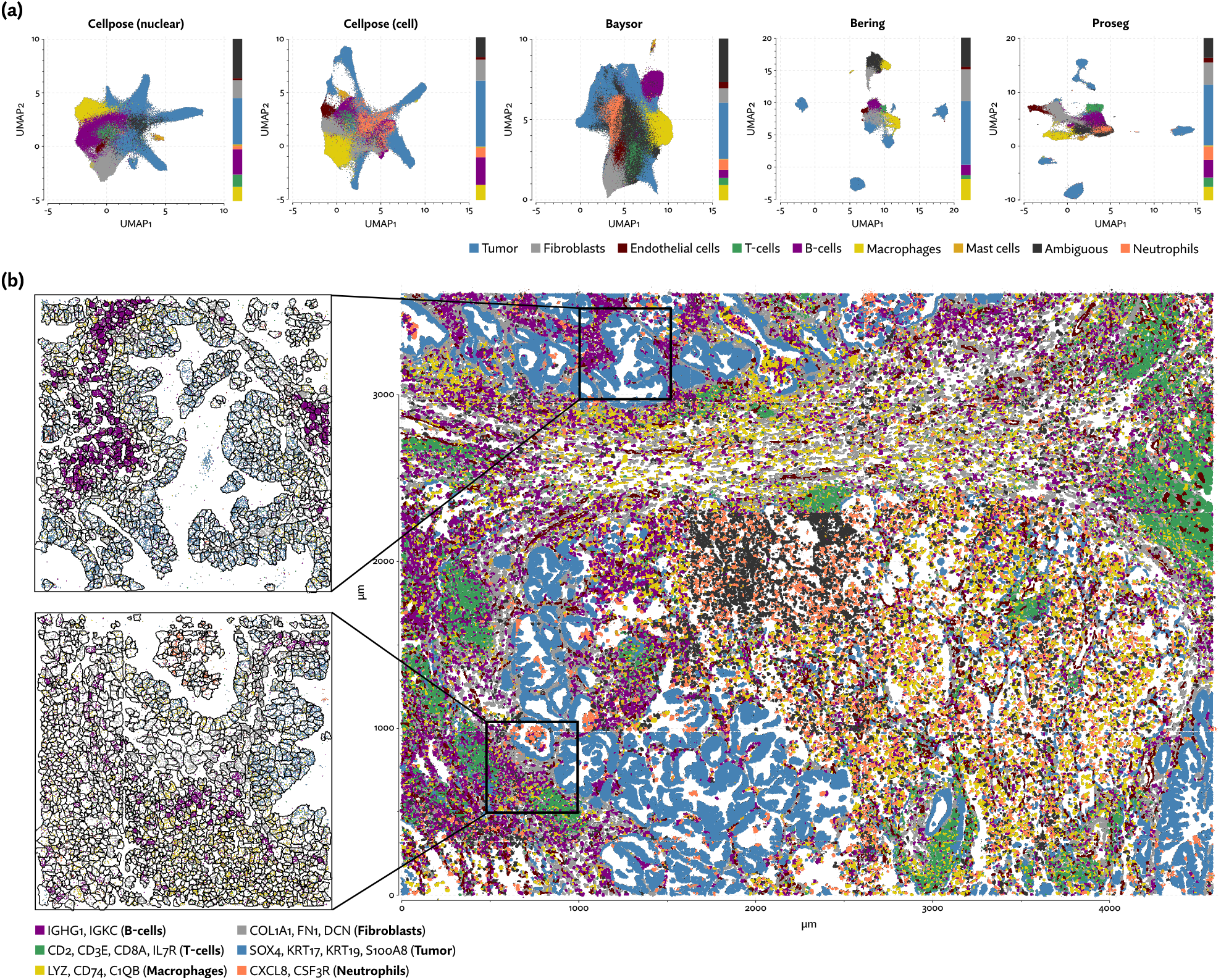
Segmentation results on the CosMx lung cancer dataset. (a) UMAP plots with annotated cell types from each segmentation method. Cell type proportions are indicated in stacked bar plots on the margins. (b) A representative example region showing cells segmented by Proseg, along with specific cell type specific sets of transcripts.

Though Bering was able to assign a large proportion of the transcripts it predicts only 131 thousand cells compared to the roughly 700 thousand predicted by Proseg and the 783 thousand predicted by the provided Cellpose model (Figures 4a, 2c). On the other extreme, Baysor predicted nearly 4 million cells. Supplementary Figure 2 shows this dramatic disagreement in one of the eight samples. While Proseg roughly agreed with Cellpose, Bering dramatically under-segmented concealing the spatial organization of the sample, and driving a very large increase in spurious co-expression (Figure 2a). Baysor, on the other hand, dramatically over-segmented causing the relatively homogenous tumor regions to appear far more infiltrated by other cell types.

Major disagreements in cell type proportion (Figure 4a, Supplementary Figure 3) were partially driven by the relative noisiness of the data. For example, major markers of T-cells and neutrophils were expressed so sparsely it was difficult to confidently label clusters in same cases. Many cells were simply labeled “ambiguous” as a result. Roughly 27% of cells segmented by Baysor are marked as such, mainly due to their extremely low transcript population.

B-cells and, once again, Neutrophils were the two cell populations with the lowest median number of transcripts, with a median of 183 and 69 transcripts per cell under Proseg, respectively, which partially explains major disagreements in their proportion. These relatively low transcript count cell types were prone to having their signal smothered by adjacent high-population cell types, specifically, fibroblasts (median of 271 transcripts per cell) and tumor cells (median of 405 transcripts per cell).

### 2.4 Reducing false-positive differential expression in tumor-adjacent macrophages in a Xenium lung cancer sample

Compared to MERSCOPE and CosMx, the initial release of 10X’s Xenium platform has relied on the less sophisticated cell segmentation strategy of nuclear expansion. Here, nuclei are identified using a deep learning model applied to a DAPI stain, then these polygons are expanded outward by some fixed distance while prohibiting overlaps. The result is a heuristic that manages to assign many of the transcripts to cells, but only in a highly approximate manner. Recently, the platform has been augmented with a full image-based cell segmentation approach using a custom deep learning model and multichannel microscopy images of nuclear, cytoplasmic, cytoskeletal, and membrane stains [10X Genomics, 2024a].

To evaluate 10X’s new segmentation approach, we downloaded a demonstration dataset generated by 10X from a human lung cancer sample [10X Genomics, 2024b]. Again, Baysor was able to remedy the spurious co-expression problem but at the cost an implausible expansion of the number of detected cells (Figure 5c), with roughly 250,000 cells compared to roughly 150,000 cells detected by the other methods (Figure 2c). Proseg was able to assign nearly all transcripts without this expansion. Cell type proportions were mostly in agreement between methods, but Baysor did diminish the number of T-cells and expanded the number of macrophages (Figure 5a). Spurious co-expression was dramatically reduced by Proseg compared to other methods on this dataset (Figure 2a).

**Figure 5:**
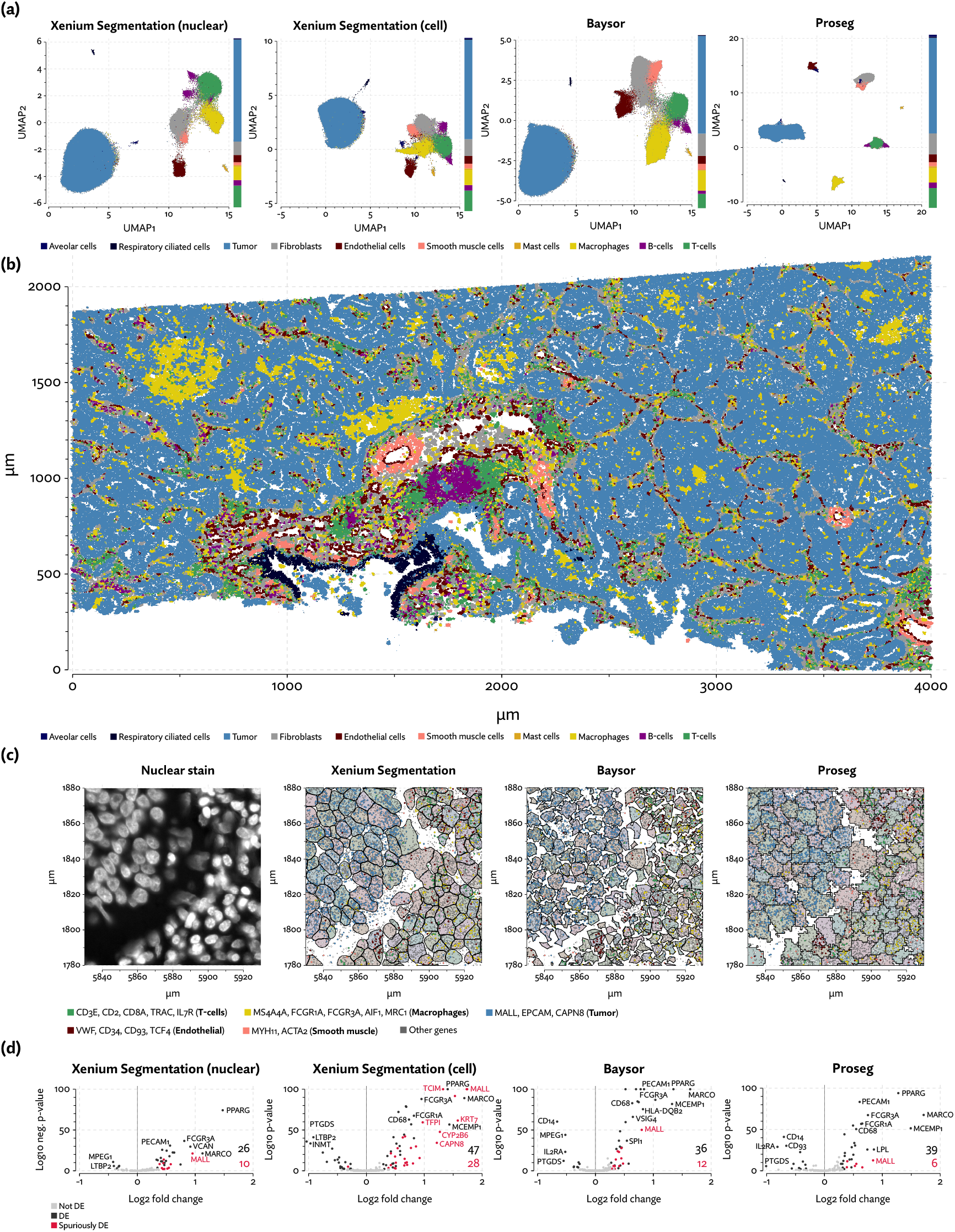
A comparison of segmentation results on the Xenium lung cancer data. (a) UMAP plots with annotated cell types from each segmentation method. Cell type composition is shown in stacked bar plots on the margins. (b) A region of the sample depicted with annotated cell types using Proseg segmentation. (c) Comparison of cell segmentation in one region, along with specific sets of highly cell type specific transcripts. (d) Differential expression results comparing tumor-adjacent to non tumor-adjacent macrophages. Labeled in red are genes that are highly expressed in tumor, thus likely spuriously called due to transcripts being misattributed to adjacent macrophages.

The sample consists of a tumor with a great deal of macrophage infiltration (Figure 5b). To demonstrate the effect of segmentation on common analyses, we performed differential expression tests using DESeq2 [Love et al., 2014] comparing macrophages adjacent to tumor cells to non-adjacent macrophages. This kind of test is easily confounded by transcripts from tumor cells being misattributed to adjacent macrophages, naively appearing as differential expression. We marked as “spurious” any differentially expressed gene with overall higher expression in tumor cells than macrophages. We cannot definitively rule out the possibility that these genes were actually upregulated in macrophages, but this pattern renders such calls suspicious at the very least. Proseg did not entirely solve this issue of suspicious differential expression calls but did greatly diminish it (Figure 5d). With the provided image-based segmentation 37% of calls were suspicious, compared to 12% with Proseg and 25% with Baysor. Using image-based segmentation with only nuclear transcripts slightly improved the situation (28% spurious calls) but reduced the statistical power of the test resulting in fewer overall calls.

Diminishing these false-positive differential expression calls helped to more accurately reveal the transcriptional program of tumor-adjacent macrophages. Macrophage heterogeneity in non-small cell lung cancer was previously explored by Casanova-Acebes et al. [2021]. This study characterized a distinct alveolar tissue resident macrophage (TRM) population. Consistent with our finding, this population was characterized by high expression of PPARG and MARCO, among other genes not present in the Xenium panel. Using fluorescent imaging they also observed the colocation of tumor cells and TRMs after initial seeding. They theorized that TRMs promote tumor invasiveness by inducing changes in epithelial cells. Though undiscussed in the paper, their data also agrees with ours in showing upregulation of MCEMP1, FCGR1A, FCGR3A, CD68, and LPL. Their analysis did not detect upgregulation of MALL, KRT7, TCIM, CAPN8, or CYP2B6, providing further evidence that these are indeed spurious calls.

### 2.5 Analysis of the tumor-proximity of CXCL13-expressing T-cells in Xenium renal cell carcinoma patient samples

To further explore the utility of Proseg, we generated Xenium data from four clear-cell renal cell carcinoma (ccRCC) patient samples each consisting of tumor and adjacent tissue (example regions shown in Figure 6b and Supplementary Figure 5). A customized gene panel was used consisting of 280 genes provided by a base panel (10X’s breast cancer gene panel) and 100 genes selected to target kidney-specific cell types, T-cell subsets, and genes that were observed to show significant variation in expression in previously generated single-cell RNA-Seq data. This data was generated prior to the recent availability of multimodal image-based segmentation for Xenium, so we compared Proseg to the nuclear expansion methodology that was provided by the platform. Nuclei were segmented using a DAPI stain, then the cell perimeter is approximated by expanding the nuclei perimeter outward by 15μm, or until another boundary was encountered (newer versions of the platform have reduced the default radius to 5μm, to lessen the missegmentation error).

**Figure 6:**
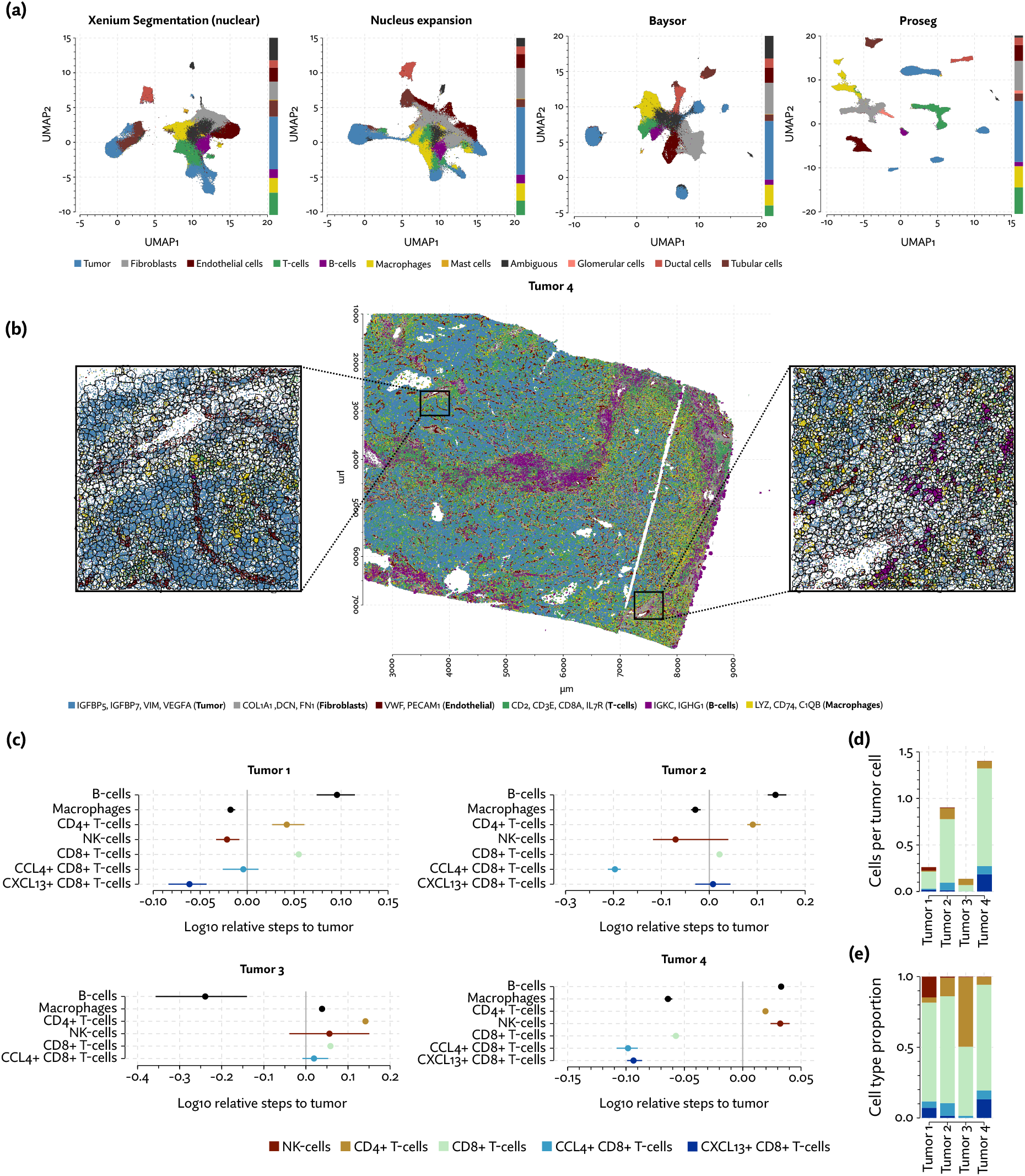
Analysis of the Xenium renal cell carcinoma dataset. (a) UMAP plots for each segmentation method, with cell type proportions shown in stacked bar plots on the margins. (b) Proseg cell segmentation in a region of one tumor sample, plotted along with subsets of cell type specific transcripts. (c) Proximity between various immune cell types and tumor cells is measured by computing the expected number of steps in a random walk on the neighborhood graph before a tumor cell is encountered, and adjusting for the local cell type composition. (d) Numbers of T-cell subtypes and NK-cells in proportion to the number of tumor cells in each tumor. (e) Relative composition of T-cell subtypes and NK-cells.

As with the other datasets, we saw the same pattern of Baysor and Proseg both reducing spurious co-expression (Figure 2c), but Baysor doing so only by introducing far more cells than other methods (Figure 2a). Both Baysor and the nuclear expansion strategy show a large reduction in the proportion of T-cells (Figure 6a, Supplementary Figure 6), consistent with them being disproportionately smothered by adjacent higher population cells. Proseg T-cells had a median of 64 transcripts per cell, far lower than, for example, macrophages (median of 180 transcripts per cell) or tumor cells (median of 226 transcripts per cell). This imbalance was roughly consistent across segmentation methods. Several of the samples show a great deal of T-cell infiltration into the tumors, exacerbating the risk of this smothering effect due to low population cells being frequently surrounded by high population cells (Supplementary Figure 7). Nuclear segmentation, which is less prone to transcript misallocation, shows a higher T-cell proportion than that seen with nuclear expansion or Baysor, consistent with the interpretation that T-cells are being concealed by missegmentation.

The degree of T-cell infiltration is generally a positive indicator of the success of immunotherapy, though this signal has proven to be far more ambiguous in ccRCC than in many other cancers [Nakano et al., 2001]. As a possible confounder, tumors can also vary dramatically in terms of the frequencies of T cells that are tumor-specific as opposed to non-specific bystanders [Simoni et al., 2018]. Numerous studies have determined that cell surface markers (e.g., CD39) and genes associated with terminal exhaustion (e.g., CXCL13) can efficiently distinguish bystander from tumor-specific CD4 and CD8 T-cells [Hanada et al., 2022, Veatch et al., 2022, Oliveira and Wu, 2023]. However, due to widespread expression of CD39 on non-T cells, quantifying and localizing CD39 expressing T-cells by multiplex immunohistochemistry, while possible, is challenging [Murakami et al., 2021, Yeong et al., 2021]. We are similarly concerned about accurately localizing T cells that do or do not express genes associated with tumor reactivity.

That the proportion of T-cells varies so dramatically between segmentation methods has concerning implications for the interpretation of the data. To further explore the significance of the T-cell populations in these samples we clustered the Proseg T-cell populations into CCL4+ CD8+ T-cells, CXCL13+ CD8+ T-cells, other CD8+ T-cells, other CD4+ T-cells, and NK-cells, the latter of which was grouped with T-cells in the broader cell type annotations. We implemented a proximity analysis based on computing the expected number of steps in a random walk on cellular neighborhood graph before a tumor cell is encountered. This number was then normalized to account for the local cell type composition (see Methods).

CXCL13+ CD8+ T-cells were primarily found in tumors 1 and 4 (Figure 6de), and in both we see a strong tendency for these cells to co-locate near tumor cells (Figure 6c). This population was not found in tumor 3 at appreciable levels, and the small number found in tumor 2 did not show a tendency to co-localize with the tumor, though CCL4+ CD8+ T-cells did in all but tumor 4. Proximity analysis can also let us measure the density of the tumor by computing the expected number of steps for a tumor cell to arrive at a non-tumor cell. From this we see that tumors 1, 4 and to a lesser degree 2 show a relatively diminished tumor density (Supplementary Figure 8), indicating a higher degree of infiltration. Broadly, in the more heavily infiltrated tumors we see a stronger tendency for one or both of the T-cell subsets to locate in the proximity of tumor cells.

Previously CXCL13+ CD8+ T-cells were found to be highly exhausted, show diminished expression of IFN-α and IFN-γ, and their intratumoral infiltration to be associated with poor prognosis [Jiao et al., 2020, Dai et al., 2021]. CCL4 can play an important but context-dependent and potentially contradictory role in the tumor microenvironment. It has been found to both contribute to recruiting tumor-promoting macrophages and regulatory T-cells, and to recruiting cytotoxic lymphocytes in different settings [Mukaida et al., 2020]. Little is known about the significance of CCL4-expressing T-cells in the context of ccRCC.

### 2.6 Discussion

Correct cell segmentation is critical to the interpretation of spatial transcriptomics data. The method proposed here has demonstrated across several datasets and platforms the ability to assign a larger proportion of the observed transcripts to a plausible number of cells, all while reducing the tendency to introduce spurious co-expression. This has clear positive effects when assigning cell types, particularly to cells with a low number of assigned transcripts, and when performing differential expression analysis. Like Baysor, by using a probabilistic model based on simple assumptions, we avoid the issue of supervision, but by using cell simulation as a starting point we arrive at a far more flexible model of cell morphology that is less prone to gerrymandering overly-parsimonious cells.

Proseg represents a very general simulation-based framework for segmenting cells. Naturally, there are numerous possible extensions and elaborations that could be pursued to improve the model. Though currently unsupervised, a straightforward extension is to allow a form of semi-supervision by pre-fitting the gene expression model to matched scRNA-Seq data. We might also improve the accuracy of segmentation by including information gleaned from image data. We do this to an extent by initializing the simulation using nuclear segmentation, and including a prior that penalizes segmentation inconsistent with the identified nuclei. The same principle can be extended by using image data to construct a more detailed prior giving voxel-level confidence in image based segmentation, either constraining or giving Proseg free rein to revise cell boundaries based on the transcript data.

An obvious limitation of the current model is that it treats cells as bags of RNA, with transcripts distributed uniformly at random throughout the cell, ignoring any subcellular localization. This idealization is close enough to reality to inform cell segmentation, but leaves room for improvement. Compartmentalized CPM models have been explored [Scianna and Preziosi, 2012, Scianna, 2015] in which nucleus, cytoplasm, and membrane have their own sub-state. Combined with an appropriate model of expression, this would allow Proseg to account for differences across cellular compartments. The second major limitation of the bag of RNA model is that we cannot count on the boundary between extremely transcriptionally similar cells to be accurately inferred. Aggregate expression estimates are largely unaffected, so this is inconsequential for many analyses, but those that rely on correct morphology may be distorted.

We have implemented model inference using the parallelization scheme proposed by Chen et al. [2007]. Though efficient, the CPM literature points towards opportunities for further gains. Tapia and D’Souza [2011] introduced a similar parallelization scheme tailored for GPUs, and more recently [Sultan et al., 2023] explored aspects of GPU implementations in detail, proposing an alternative scheme that is faster while avoiding potential inconsistencies introduced with the earlier scheme, accelerating performance by orders of magnitude over a CPU implementation.

Cell segmentation is, and continues to be, the biggest hurdle when attempting to use in situ spatial transcriptomics to study the relationship between gene expression and spatial organization. More work will be required before we can claim that the problem is truly solved, but with Proseg we have introduced a method that has dramatically reduced the confounding of expression and proximity taking a major step towards trustworthy analysis.

## 3 Methods

### 3.1 Representation of data and cell boundaries

A spatial transcriptomics dataset 𝒯 = (g, s) is assumed to consist of M RNA transcripts from a gene panel of m genes. Each transcript i has a gene g_i_ ∈ {1, …, m} and spatial coordinates *s*_*i*_ ∈ ℝ^3^. The goal of cell segmentation is to infer which of these transcripts is background noise (due to optical shot noise or extracellular RNA, for example), and partition the rest into cells. Proseg, like most segmentation methods, does this by determining non-overlapping cell boundaries, and attributing to a cell any transcript in within its boundary that is not inferred to be background noise.

We make the simplifying assumption that cell nuclei can be roughly identified from a DAPI stain. We use this information to constrain the number of cells and penalize nuclear transcripts being assigned to another cell. This initial nuclei segmentation is unlikely to be perfect, but initializing and informing the model with these landmarks simplifies and speeds up inference, and lessens the risk of over-segmenting. Relying purely on a model of gene expression otherwise risks gerrymandering cells into implausibly parsimonious cell fragments.

Broadly, we model the spatial pattern of transcript expression under a simple Poisson point process. Where the Poisson rate parameter for each gene is uniform across a cell’s volume and governed across cells by a Gamma mixture model. The boundary of a cell’s volume is inferred by discretizing space into voxels and sampling the assignment of voxels to cells according to this probabilistic model of gene expression, with constraints on its connectivity and surface area. The assumption that transcripts of a gene are uniformly distributed within a cell is of course not strictly true — mRNA can be localized to particular subcellular regions, or never exported from the nucleus in the case of some noncoding RNA — but true enough in principle to inform segmentation.

### 3.2 Sampling cell boundaries

Much of the inspiration and methodology for Proseg comes from Cellular Potts models (CPMs). These models are a probabilistic variant of cellular automata that seeks to induce biologically plausible cell behavior from simple rules governing the energy state of the system. Briefly, a population of cells are simulated on a (typically) rectangular lattice of 2-dimensional pixels or 3-dimensional voxels. Each of these voxels is either unassigned or assigned to exactly one of the cells.

The CPM sampling strategy is to generate random update proposals, each proposal consisting of copying the state of one voxel i to a neighboring voxel j. We abbreviate these state change proposals as i→j. CPMs then accept or reject these proposals using a simulated annealing optimization scheme. Specifically, the change in energy ΔH(i → j) is evaluated, and the proposal is accepted with probability

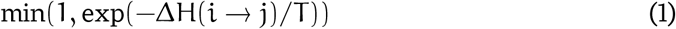

where T is a temperature parameter, which is either fixed at some reasonable value or annealed towards zero during optimization. How energy is modeled governs what cell-like behaviors the system produces. Much of CPM research has focused on constructing energy functions that reproduce observed real world behaviors such as cell type sorting [Graner and Glazier, 1992] and migration [Niculescu et al., 2015].

Our approach to segmentation borrows its basic representation and sampling procedure from CPMs, but transforms it into a Bayesian probabilistic model of the observed data, accepting or rejecting random proposals according a Metropolis-Hastings algorithm. This lets us simulate cells that conform to the underlying data (i.e., the observed transcripts), rather than just an ab initio energy function, and lets us potentially quantify the uncertainty of cell boundaries and transcript assignments.

To formalize this idea, each voxel’s state is either unassigned, which we denote as ∅, or assigned to one of n cells (i.e., in {1, …, n}). The state of the model is then denoted 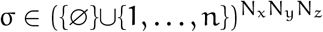, where N_x_, N_y_, N_z_ are the number of voxels in each respective dimension.

Let *σ*_*i*_ ∈ {∅}∪{1, …, n} be the current state assignment of voxel i, and σ_i→j_ the state formed by taking σ and copying the value in voxel i to voxel j. The posterior probability of a state σ, conditioned on the observed transcripts 𝒯 and model parameters θ, is denoted P(σ|𝒯, θ). The details of this model will be described in a following section.

Under to Metropolis-Hastings rule, we then accept the *i* → *j* proposal with probability

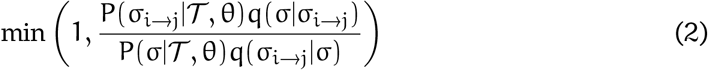

in which q is the proposal distribution, where q(σ ^*′*^|σ) gives the probability of a transition from state σ to σ ^*′*^being proposed.

This closely resembles the CPM update rule (Equation 1), but substituting an explicit probability distribution in place of the (negative exponential) energy function. There are two other notable changes. First there is no temperature parameter, or equivalently T = 1. Second, we must weight acceptance probabilities by the proposal distribution q to preserve detailed balance, ensuring the sampler correctly draws from the posterior probability distributionneighboring

To generate state change proposals we maintain a set of every pair of voxels (*i, j*) in the same von Neumann neighborhood (i.e., a cell and its 6 neighbors that share a voxel face) in which σ_i_ *≠* σ_j_. Edges are selected uniformly at random from this set on each iteration.

### 3.3 Ensuring detailed balance

The correctness of any Markov-chain Monte Carlo (MCMC) scheme rests on the preservation of detailed balance. Lacking this, there is no guarantee of long-run convergence to the stationary distribution, and no basis for treating samples as arising from the posterior probability distribution. This is less of a concern for traditional CPMs, which seek to optimize the energy function rather than sample from a posterior probability distribution.

For Metropolis-Hastings samplers, the primary concern is that every state change proposed is reversible with an inverse proposal. Concretely, if a state change *i* → *j* is irreversible, then q(σ|σ_i→j_) = 0, and the acceptance probability in Equation 2 is also 0, closing off a possible path for the sampler to explore.

A naive implementation of a CPM inspired sampling scheme, consisting only of propagating neighboring states, does not obey the detailed balance condition. Violation occurs by generating irreversible proposals, such as those that annihilate a cell completely (Supplementary Figure 9a) with no way for it to be reborn, or those that pop bubbles that then cannot reform (Supplementary Figure 9c,e,g). As a result, we have to take care to either avoid making irreversible proposals, or to modify the procedure to make some naively irreversible proposals reversible.

The irreversibility of cell annihilation is easily addressed by prohibition: if a cell is reduced to a single voxel, we do not make proposals that would change its state. Much more difficult is the issue of popping bubbles. Our scheme borrows an approach proposed by Durand and Guesnet [2016] for CPMs. The authors show that preserving the local connectivity of each cell is a sufficient condition to guarantee detailed balance.

A state change to a voxel j modifies the local connectivity if it changes the number of connected components of any cell within the Moore neighborhood of j (i.e., j and its 26 bordering voxels). On a proposal i→j we check if j in an articulation point for either cell σ_i_ or σ_j_ within j’s Moore neighborhood. It is an articulation point if and only if changing its assignment modifies the number of local connected components. We pre-reject the proposals without evaluating their probability, ensuring we avoid an irreversibility state changes.

We make an exception for the ∅ state. Preserving local connectivity of unassigned regions would preserve detailed balance, but would be severely limit the ability of the sampler to explore biologically plausible states. Most significantly, we could not form nor pop inter-cell ∅ bubbles (Supplementary Figure 9f,g). We instead allow local connectivity changes of the ∅ state and deal with bubble popping by making it a reversible state change.

To do so, we introduce random ab nihilo ∅ bubble formation proposals (Supplementary Figure 9h). Random i→j proposals, with some small probability (we use 0.05 in practice) will update j with ∅ rather than σ_i_. This lets ∅ bubbles form and pop between cells while maintaining detailed balanced.

### 3.4 Constraining perimeter to volume ratios

Left to its own devices, CPMs and CPM-inspired segmentation can produce cells with implausible morphology. As mentioned, the primary risk in driving cell segmentation by modeling of gene expression is that cells are gerrymandered into tortured shapes to induce a higher probability state. We avoid this situation by constraining each cell’s surface area to volume ratio. Doing so, we enforce some level of compactness while avoiding making any specific morphology assumptions.

In imaging based spatial transcriptomics, x- and y-coordinates of transcripts are fairly precise, while the z-coordinate, if provided, is based on focal length, so cannot be assumed to be as precise. Thus, segmentation on the z-axis is more approximate, and in practice we use lower resolution in this dimension. In keeping with this quasi-3D setting, we constrain the perimeter of each cell separately on each layer of voxels on the z-axis.

Taking one such layer belonging to a cell, we define the volume as the number of voxels making up that layer of the cell.

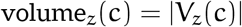

The perimeter of that layer is then the number of bordering neighbors not in the cell.

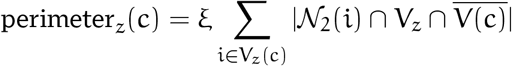

where 𝒩_2_(i) is the Moore neighborhood of voxel i. As explored by Magno et al. [2015], counting neighbors this way is approximately proportional to the *perceived perimeter*, roughly meaning the perimeter after smoothing the voxel region, after scaling by an appropriate constant ξ.

The perimeter of a circle with area a is 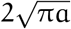. We constrain the perimeter of a cell at layer z to be a factor b of this.

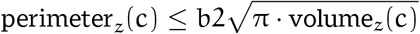

In practice, we use b = 1.3 by default. A sampler proposal that exceeds this is automatically rejected. Some cell types have far more complicated morphology, so relaxing the parameter could be appropriate in those settings, provided the data is high enough resolution to resolve the true morphology.

### 3.5 Sampling voxel updates in parallel

At high resolutions, a region make be composed of a very large number of voxels. For efficiency reasons, we would like to make many state change proposals in parallel, which both reduces overhead and allows parallelization across many CPU cores.

In general, parallel updates are only possible by exploiting a conditional independence structures in the model. In our model, every transcript is either background (∅ state) or belongs to a cell whose boundary it resides in, where cell boundaries are strictly disjoint. Because of this, a change to a cell’s boundary affects the likelihood only of the transcripts within its boundary. Evaluating a proposal i→j necessitates recomputing only the likelihood of the transcripts in cells σ_i_ and σ_j_, assuming neither is ∅. We are therefore free to carry out parallel updates so long as no two updates affect the same cell.

We adopt a simple procedure to ensure this, which was proposed in CPMs by Chen et al. [2007]. The region is divided into bins, then each bin further subdivided into four sub-quadrants. The sampler proceeds in four phases, making parallel proposals on every first, second, third, and fourth sub-quadrant, before cycling back around the first (Supplementary Figure 10). If the size of the quadrants is chosen to be sufficiently large, then the gap between each kth quadrant ensures that no two proposals involve the same cells.

Independent updates are not strictly guaranteed by this procedure; nothing prevents a cell from growing large enough to span a quadrant and become involved in multiple updates. A small nonzero number of inconsistent parallel updates being made is not catastrophic, but in practice we can calibrate the quadrant size using known nuclei positions to ensure this almost certainly does not happen. Though not yet implemented, this parallelization scheme also opens the possibility of even faster GPU implementation, as has been done with CPMs [Tapia and D’Souza, 2011].

### 3.6 Varying resolution

Partitioning space into voxels necessitates choosing a resolution, or equivalently the dimensions of the voxels. Naively it would seem that the higher the resolution the better. Though it does lead to more precise cell boundaries, a higher resolution can result in the sampler taking many iterations to work its way out of a lower probability state.

To overcome this trade-off, we begin sampling with relatively large voxels, then periodically double the resolution during sampling. Doubling resolution splits each voxel into eight sub-voxels, but does not alter the region assigned to any cell, letting the sampler move forward proposing more detailed changes. In practice, we initialize with voxels that are 4μm on the x and y axis, and a single layer on the z axis (encompassing the span of z coordinates). We then use a sampling schedule that halves these dimensions twice during sampling ultimately arriving at 1μm voxels, stacked 4 layers high on the z axis.

### 3.7 Modeling a cellular gene expression

Reiterating the broad representation used, the region under consideration is discretized into layers of voxels, which may vary in dimensionality. Each voxel i has a corresponding state σ_i_ ∈ {∅} ∪ {1, …, n}, either assigning it to a cell or leaving it unassigned (which we denote as ∅).

To describe occurrences of transcripts in and outside of cells, we adopt a Poisson point process model. The concentration of transcripts of each gene g is governed by a rate function λ_g_ that varies across voxels. This function is the sum of two parts: the background rate (which may be due to optical shot-noise, nonspecific probe binding, or ambient extra-cellular RNA), and the cell rate, describing the contribution of any overlapping cell.

We define the background rate to be constant, though one could imagine a more elaborate model that tries to account for possible covariates like field of view or broad spatial context. For voxel i, we define the background rate as

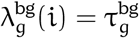

Similarly, cell rates are also defined as constant on a per-cell basis

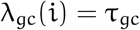

For convenience we define this for the unassigned ∅ state as well.

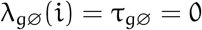

Now, the rate for any gene g and voxel i can be written

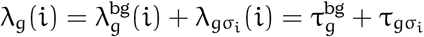

We can then write the Poisson point process likelihood as

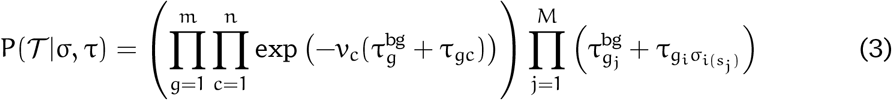

where v_c_ is the volume of cell c, which is simply the number of voxels |V(c)| scaled by the volume of each voxel, and i(s_j_) is the voxel containing the position s_j_ of transcript j.

As noted, a critical feature of this likelihood function is that accepting an update proposal i → j only affects factors in the likelihood corresponding to the effected cell or cells, and the transcripts in those cells. This lets us sample cell boundaries in parallel so long as we can avoid simultaneous changes to the same cell.

We model cell-rates according to hierarchical Gamma mixture model with k components (Supplementary Figure 11). The intuition here is that mixture model components represent broad cell types. It is not essential to the model that the number of components precisely match the number of “true” cell types. Indeed, the model can be run with a single component, but representing cells as belonging to different components helps the model distinguish between transcriptionally distinct types. More explicitly, the mixture model is defined as follows.

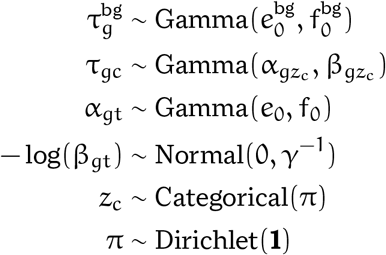

In words, the rate τ_gc_ for each gene and cell (interpretable as the expected gene expression, normalized by cell volume), is Gamma distributed, according to the cells component z_c_. The choice of Gamma distributed gene expression rates means that cell gene counts are Gamma-Poisson, or equivalently negative binomial distributed. This lets us efficiently sample certain posterior parameters.

We make two other minor additions to the expression model. First, we model the volume of cells by component, to potentially capture major differences in cell sizes.

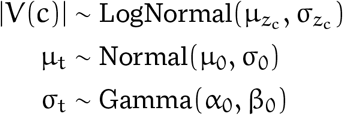

Secondly, we use a prior probability of the correctness of the initial assignment of transcripts to identified nuclei. Where the initial state is 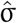, we include a nuclear prior term in the model,

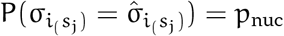

where we use p_nuc_ = 0.8 in practice, penalizing reassignment.

### 3.8 Modeling the phenomenon of leaky cells

A standard assumption made by cell segmentation algorithms is that observed RNA all reside within their originating cell with distinct boundaries. Partitioning space is thus, for most purposes, interchangeable with partitioning transcripts. If we can determine the precise boundaries of a cell in space, then a transcript belongs to that cell if and only if it resides within those boundaries.

What we observe, particularly in some Xenium data, is a tendency for transcripts to leak from cells, and diffuse to nearby regions. These are frequently at a lower focal length than transcripts within the tissue (Supplementary Figure 12a), indicating the transcripts are on the slide surface, having diffused off the sample. Furthermore, the gene composition of transcripts outside the tissue region very closely matches that of transcripts in the tissue (Supplementary Figure 12b). This confounds downstream analysis, introducing cross-talk between neighboring cells that no segmentation algorithm based on the standard assumption can hope to resolve.

Because Proseg is a probabilistic model, we can include in a principled way the possibility of transcripts being diffused from their originating cells. We do this by placing a prior on the distance a transcript migrates from its true position and, in addition to sampling cell boundaries, we sample transcript repositioning.

Where 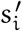 is the observed position of transcript i, we posit a true position *s*_*i*_ with Normal mixture prior

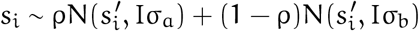

In practice, we set ρ = 0.2, σ_a_ = 4, σ_b_ = 0.5. Ideally these parameters would be modeled as well, but such a model is difficult to fit. Without any constraining assumptions, it can easily work its way into a state where everything is assumed to be widely diffused.

### 3.9 Parameter inference

As we are assuming the distribution of non-background transcripts follows a Poisson point process, thus the number of transcripts of gene g in cell c follows a Poisson distribution

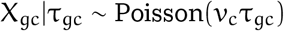

The per-volume rate τ_gc_ has a Gamma prior. A scaled gamma random variable is itself gamma, so we have,

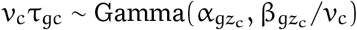

Thus, the marginal distribution of X_gc_ is negative binomial

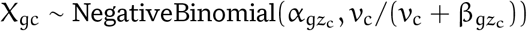

#### Sampling component assignments

Exploiting the negative binomial marginal, we can efficiently sample component assignments

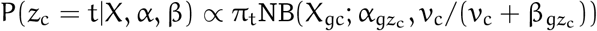

where NB is the negative binomial probability mass function.

#### Sampling Poisson rates

Exploiting Gamma-Poisson conjugacy, we can sample the rate parameter for cell c and gene g as

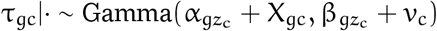

#### Sampling α

The inference scheme for the α and β parameters borrows from the model of scRNA-Seq developed by Dadaneh et al. [2020]. The α parameters are sampled using a data augmentation scheme derived by Zhou and Carin [2015] based on the Chinese restaurant table (CRT) distribution, written

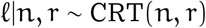

which can be generated by summing Bernoulli distributed variables

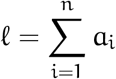

where a_i_ ∼ Bernoulli 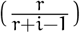.

CRT distributed random variables are drawn for each gene and cell pair,

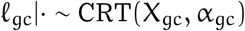

The posterior for α_gt_ is then Gamma distributed as,

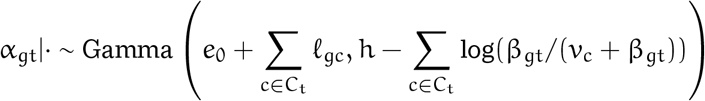

where C_t_ = {c ∈{1, …, n}|z_c_ = t} is the set of all cells in component t.

The posterior of the h parameter, which influences the overall dispersion of components, is Gamma distributed as,

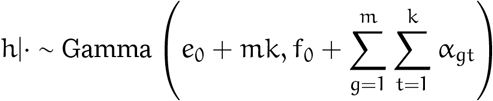

#### Sampling β

The β parameters are sampled using a Polya-Gamma augmentation scheme developed by Zhou et al. [2012] and Polson et al. [2013].

For convenience, we introduce a reparameterization of β:

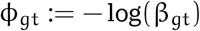

We then consider the logit transformation of the negative binomial p parameter,

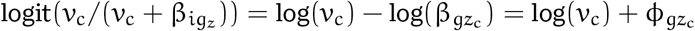

Auxiliary Polya-Gamma random variables are drawn for each gene and cell pair

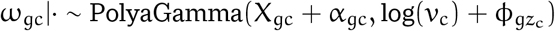

for g ∈ 1, …, m and c ∈ 1, …, n.

The posterior of 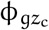 is then

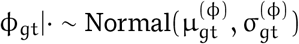

Where

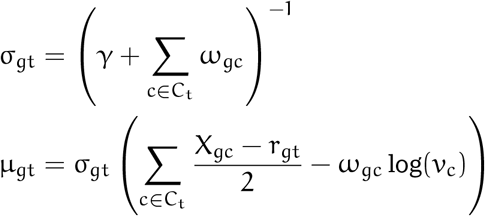

And β is derived from the reparameterization as β_gt_ = exp(−ϕ_gt_).

### 3.10 Mathematical notation reference

*m* Number of genes

*M* Number of observed transcripts

*n* Number of cells

*N* Number of voxels

*b* Spatial volume of one voxel

N_x_, N_y_, N_z_ Number of voxels in the x, y, z dimension, resp.

*k* Number of components

*g*_*j*_ Gene of transcript j in {1, …, m}.

*s*_*j*_ Spatial coordinates of transcript j in ℝ ^3^

*σ*_*i*_ State (cell-assignment) for voxel i

*i*(*s*) The voxel containing point s ∈ ℝ^3^

∅ A voxel state indicating it is unassigned to any cell

V Set of total voxels composing the region being considered

V(c) Set of voxels assigned to cell c (equiv. {i *∈* V|A_i_ = c})

V_z_ ⊂ V Set of total voxels in a particular z-layer

V_z_(c) V_z_ ∩ V(c)

v_c_ := b|V(c)| Spatial volume of cell c

𝒩_1_(i) The von Neumann neighborhood of voxel i

𝒩_2_(i) The Moore neighborhood of voxel i

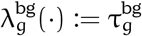Poisson background rate over voxels for gene g.

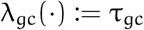 Poisson rate over voxels for gene g and cell c.

z_c_ Component of cell c in {1, …, k}.

C_t_ Set of cells belonging to component t (equiv. {c *∈* {1, …, n}|z_c_ = t}) α_g_ Gamma shape parameter for gene g

β_gt_ Gamma rate parameter for gene g and component t

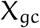 Number of non-background transcripts of gene g in cell c

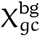 Number of background transcripts of gene g in cell c

γ Precision parameter for the prior on β_gt_

### 3.11 Data acquisition and processing

#### 3.11.1 Xenium data acquisition

The RCC FFPE blocks were provided by Northwest Biotrust, under a NWBiospecimens protocol, Seattle. NW BioTrust, a core service for patient consenting, and NWBioSpecimen, a core service for procurement and annotation of research biospecimens. The analysis was performed according to the IRB file/approval number NHS #6007-1061. No age or gender information is available. Informed written consent was obtained from each subject or each subject’s guardian.

Xenium allows customization of gene panels, which we took advantage of for this project. We selected 100 genes to add on to a standard breast cancer base panel of 280 genes. Fourteen of these were manually selected from prior interest and the remaining 86 were selected using an existing in an unsupervised manner using a prior scRNA-Seq dataset. Briefly, multinomial logistic regression was fit to cell type labels in this data and genes were prioritized by their absolute regression coefficient, as an approximation of how useful the gene is in distinguishing known cell types. The 100 add on genes were finalized in collaboration with 10X, removing some with very high expression risking optical crowding.

We followed the Demonstrated Protocol Xenium in Situ Tissue Preparation Guide (CG000578 Rev A) to place samples on Xenium slides (10.45mm x 22.45mm). Deparaffinization and decrosslinking steps followed protocol CG000580 (Rev A). The Xenium In Situ Gene Expression User Guide (GC000582 Rev A) was used for the remaining workflow. The human breast base panel (280 genes) and a 100 gene custom add-on panel (See supplementary) were combined at the probe hybridization step. The slides were loaded in the Xenium Analyzer as directed by CG000584 (Rev A). Imaging was conducted in cycles on the instrument, capturing data across multiple Z-planes and fluorescence channels to build a spatial map of transcripts within selected scanned regions. For two samples we selected five regions within each tissue representing tumor and adjacent based on histology. We scanned the entire tissue as a single region for the second pair samples. The onboard analysis pipeline included image processing, nucleus and cell segmentation, RCA product image registration, transcript decoding, quality scoring, and deduplication. Quality scores were estimated using controls for calibration, ensuring the accuracy of the final output data. After decoding, duplicate transcripts were reconciled, and a cell-feature matrix was generated, allowing for further analysis and integration with existing datasets.

#### 3.11.2 Data processing

We followed a similar strategy when processing each spatial transcriptomics dataset presented. Each segmentation method ultimately produces a cell-by-gene matrix of transcript counts, and at least spatial coordinates of each cell’s centroid, if not full cell boundary coordinates. We first excluded cells with fewer than 10 transcripts.

We then generated dimensionality reduction using UMAP and unsupervised clustering using Leiden. The same k-nearest neighbors graph was used for both, with k = 15 and nearness defined using Euclidean distance applied to censored log-proportions. Specifically, each cell’s count vector is divided by its sum, bounded below by 10^−4^, and log transformed before computing distances. Proportions potentially discard some information (indeed, differences between cell type average transcript counts is a major theme here), yet are less prone to introducing batch effects between samples or runs, so are preferable in this setting.

#### 3.11.3 Testing for spurious co-expression

To quantitatively assess the quality of segmentation we designed a benchmark to approximate the degree of spurious co-expression due to mis-segmentation. Our starting assumption is that nuclear segmentation, being an overall easier segmentation problem, will have a lower rate of incorrectly assigned transcripts than full-cell segmentation. Thus, if the rate of co-expression is dramatically higher in full-cell segmentation, it’s likely due to missegmentation. This is an imperfect assumption because this increase in co-expression could also be explained by subcellular location to the cytoplasm, but presence in the cytoplasm and absence from the nucleus is a somewhat rare phenomenon.

We first identified a set of genes that will appear spuriously co-expressed under low-quality segmentation. To do this we compared nuclear segmentation to intentionally low-quality nuclear expansion segmentation. Let P_i_ denote the set of cells expressing gene I (that is, with count ≥ 1). We define the *conditional co-expression* between genes i and j as

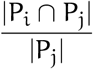

Let C_ij_ give the conditional co-expression score under nuclear segmentation for genes i and j, and 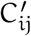 the same under nuclear expansion segmentation. We define as spurious any pair of genes (i, j) where 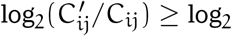.

To produce a relative spurious co-expression distribution, we computed the conditional co-expression for each method and each pair of spuriously co-expressed genes, then normalized (i.e., divided) by the rate observed under nuclear segmentation.

This metric can be cheated to some extent by either assigning fewer transcripts, or predicting an implausibly large number of cells. If less is co-expressed, there is less spurious co-expression. To counter this, we used a down-sampled counts by selecting just cells with at least 50 transcripts, then sampling a new count vector from a Multinomial distribution with a total count of 50 and probabilities matching the observed proportions for that cell. As a result, spurious co-expression is assessed on data where each cell has an exact count of 50 transcripts, but matches the observed proportions in expectation.

#### 3.11.4 Measuring cell proximity

To measure the overall proximity of a cell to cells of a particular type, we borrow concepts from Markov chain analysis. On a neighborhood graph between cells we consider a random walk process across neighbor edges in which cells of a particular type — in our case, tumor cells — are treated as “absorbing”, lacking any outgoing edges. The walk terminates when one of these absorbing cells in encountered. An absorbing Markov chain can be described by a transition probability matrix of the form

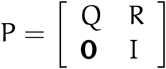

where the sub-matrix Q is the transition probability between non-absorbing *transient* cells, and the sub-matrix R the transition probabilities from transient to absorbing cells.

We measure the overall proximity from each transient cell to tumor cells by considering the expected number of steps on this random walk before absorption. This is also referred to as the *expected hitting time*, which can be defined as

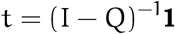

This can be computed efficiently by solving the following sparse linear system using any off-the-shelf solver

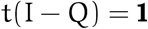

When *t*_*i*_ is small, then cell i is considered to be in close proximity to tumor cells. Importantly, this measure is not a distance, being inherently non-symmetric. For example, macrophages being near tumor cells on average does not imply tumor cells are near macrophages on average.

We define the transition probability matrix using the Delaunay triangulation, deleting edges outgoing from absorbing states and assigning uniform probability to outgoing edges. The method is agnostic to how the neighborhood transition matrix is defined, so it might be elaborated on by weighting by inter-cell distance, or using a k-nearest neighbor graph.

Aggregate proximity is heavily influenced by the local cell type composition which complicated meaningful comparison. The random walk methodology adapted to effectively normalize out this influence to get directly at the question of co-localization independent of local composition.

To produce a normalized expected hitting time, we first send each cell on a k step random walk on the Delaunay graph before measuring expected hitting time. This can be thought of as shuffling within a local neighborhood, preserving local cell type composition but not the specific organization. This process is repeated and averaged to estimate the background expected hitting time. The relative expected hitting time is then ratio between the measured and background expected hitting times. This measure is then 1 when proximity is what it would be under random organization of the same cells, and more or less than 1 to the degree that the specific cellular organization in non-random.

## 4 Data and code availability

Proseg is available under and open source license at https://github.com/dcjones/proseg. The Xeinum RCC transcript data is available at DOI: 10.6084/m9.figshare.25685961.

