## Supplemental Figures for "Cell Simulation as Cell Segmentation"

### Cell Simulation as Cell Segmentation: Supplementary Figures

July 2, 2024

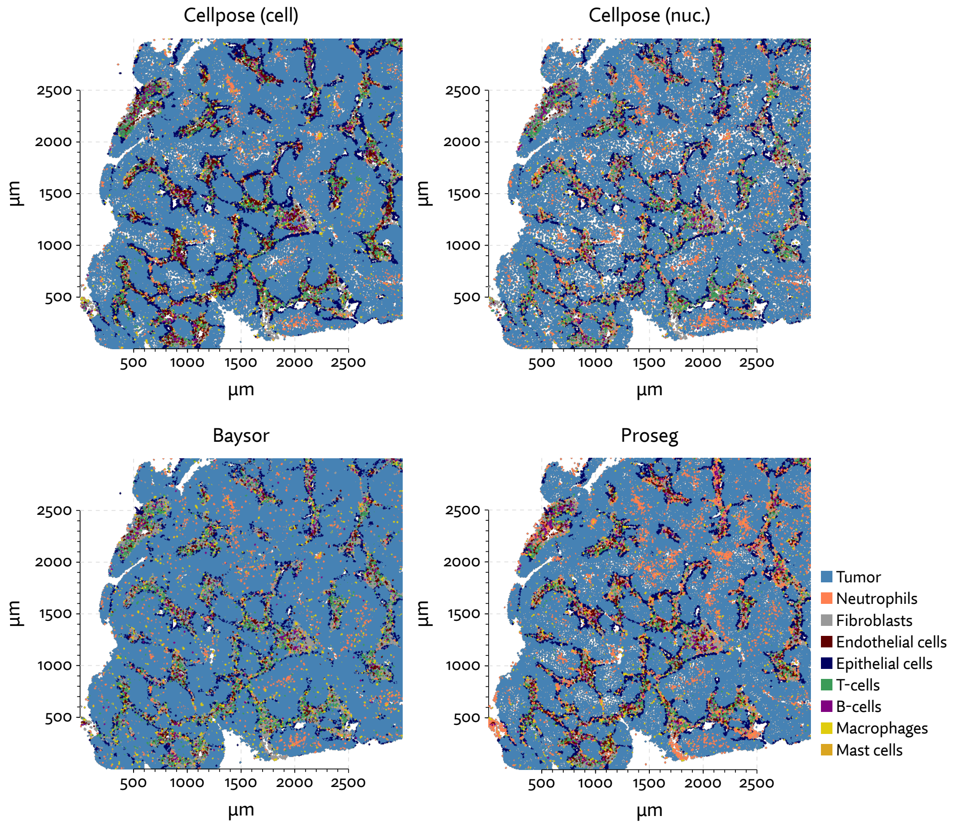


**Supplementary Figure 1:** A comparison of the MERSCOPE lung cancer dataset segmented using the default Cellpose approach and with Proseg showing the undercounting of neutrophils.

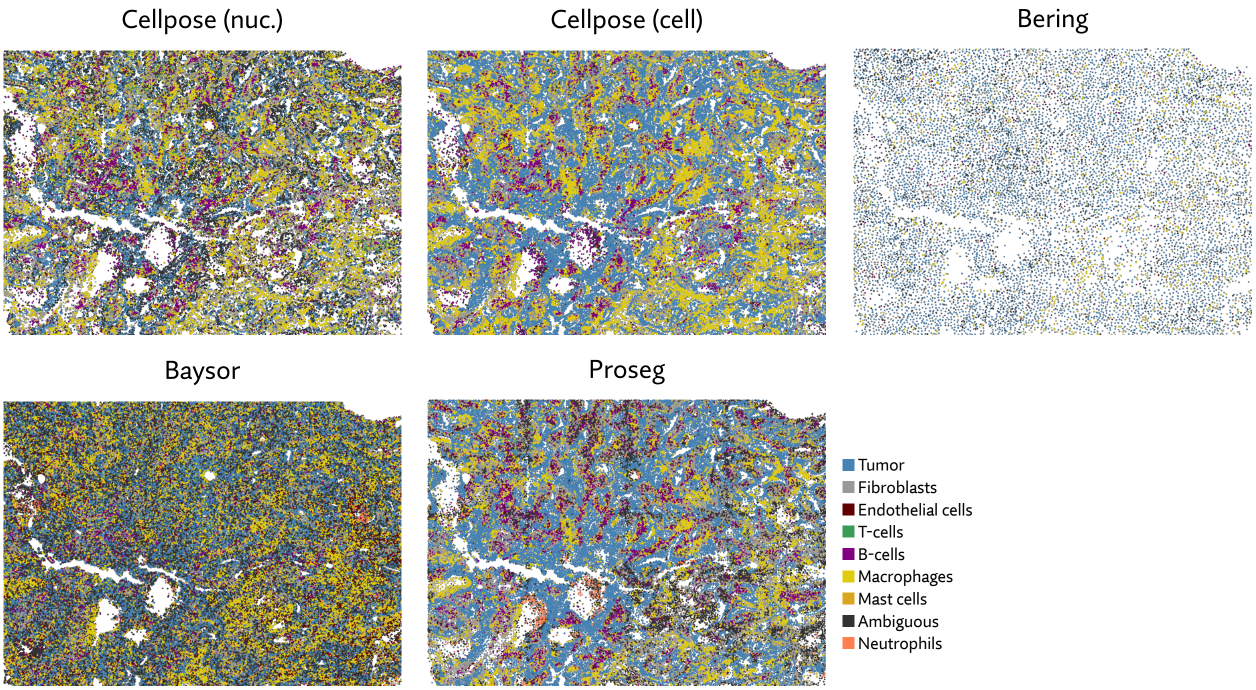


**Supplementary Figure 2:** Annotated cell types across each segmentation method on a selected CosMx sample. Segmentation methods differ dramatically on the number of cells and apparent cell types.

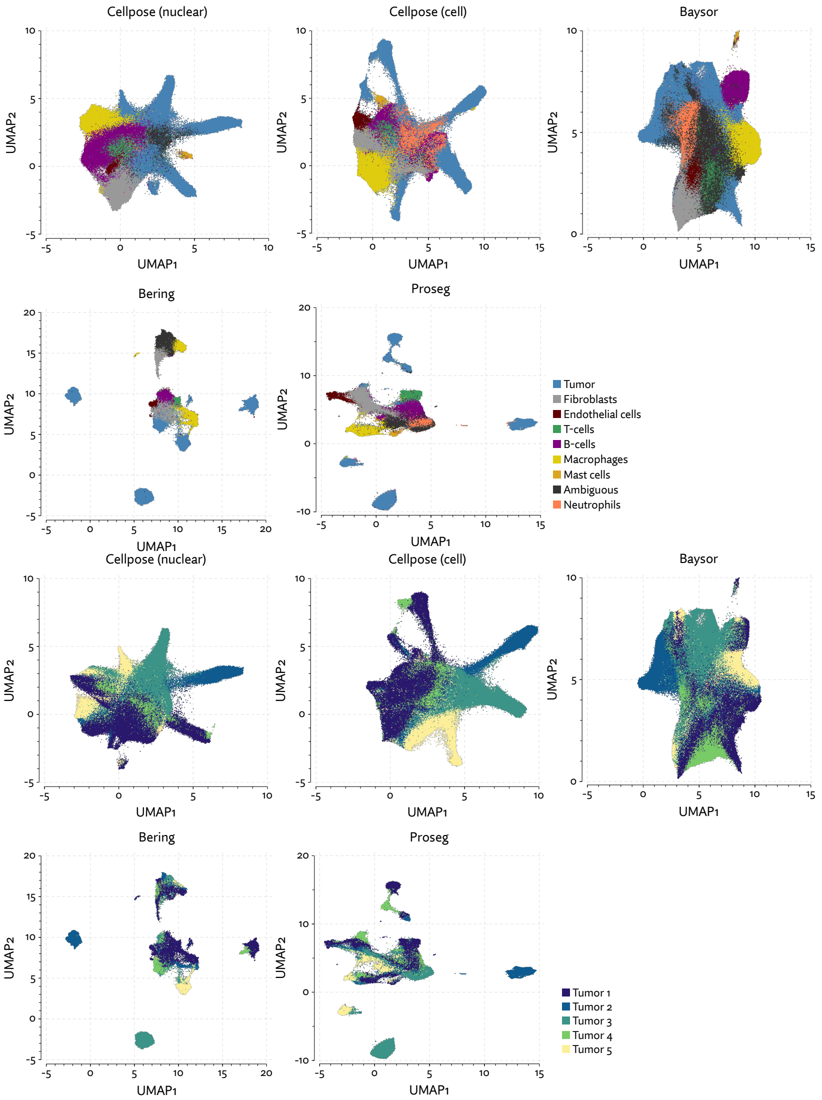


**Supplementary Figure 3:** UMAP plots for the CosMx lung cancer dataset using different segmentation methods.

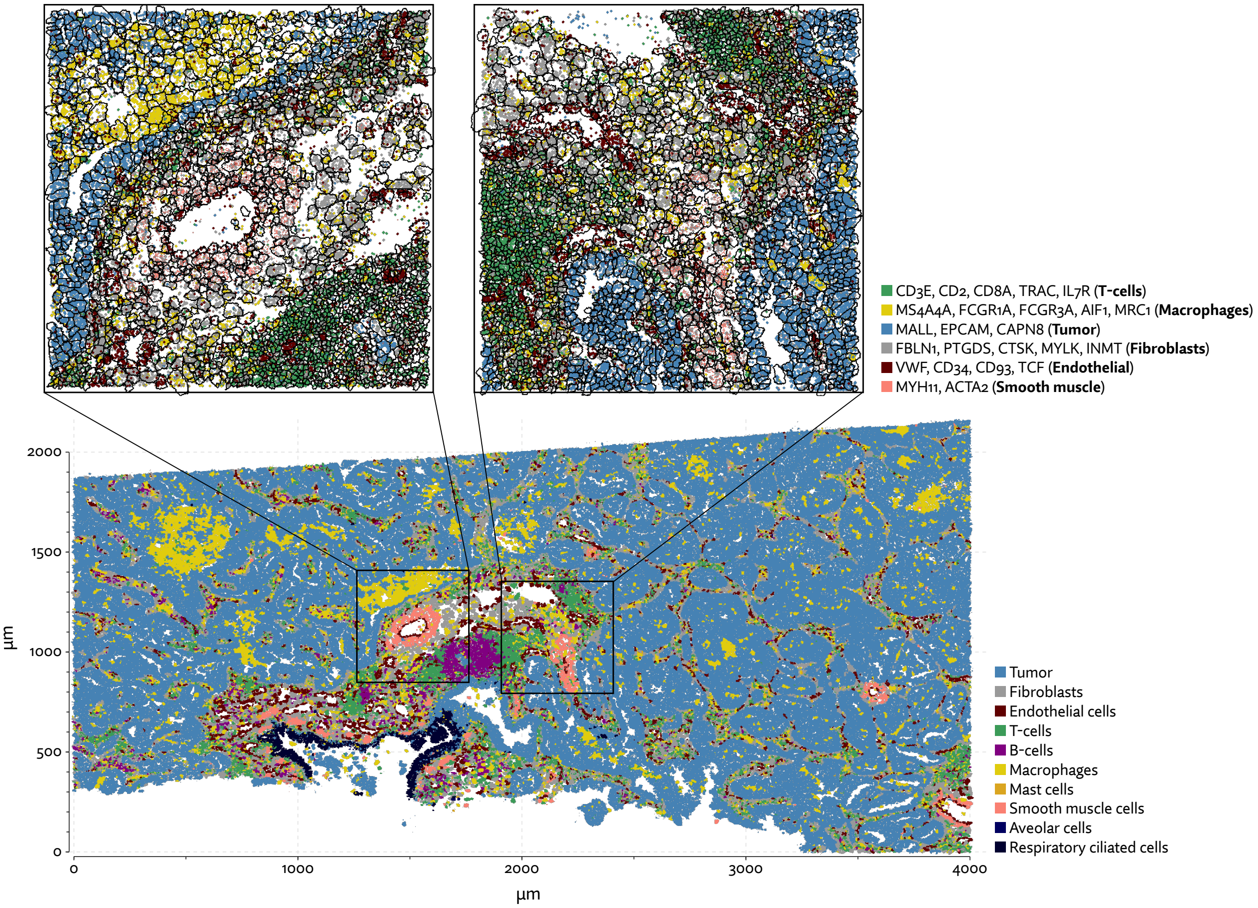


**Supplementary Figure 4:** One region of the Xenium lung cancer dataset with annotated cell types after segmenting cells with Proseg.

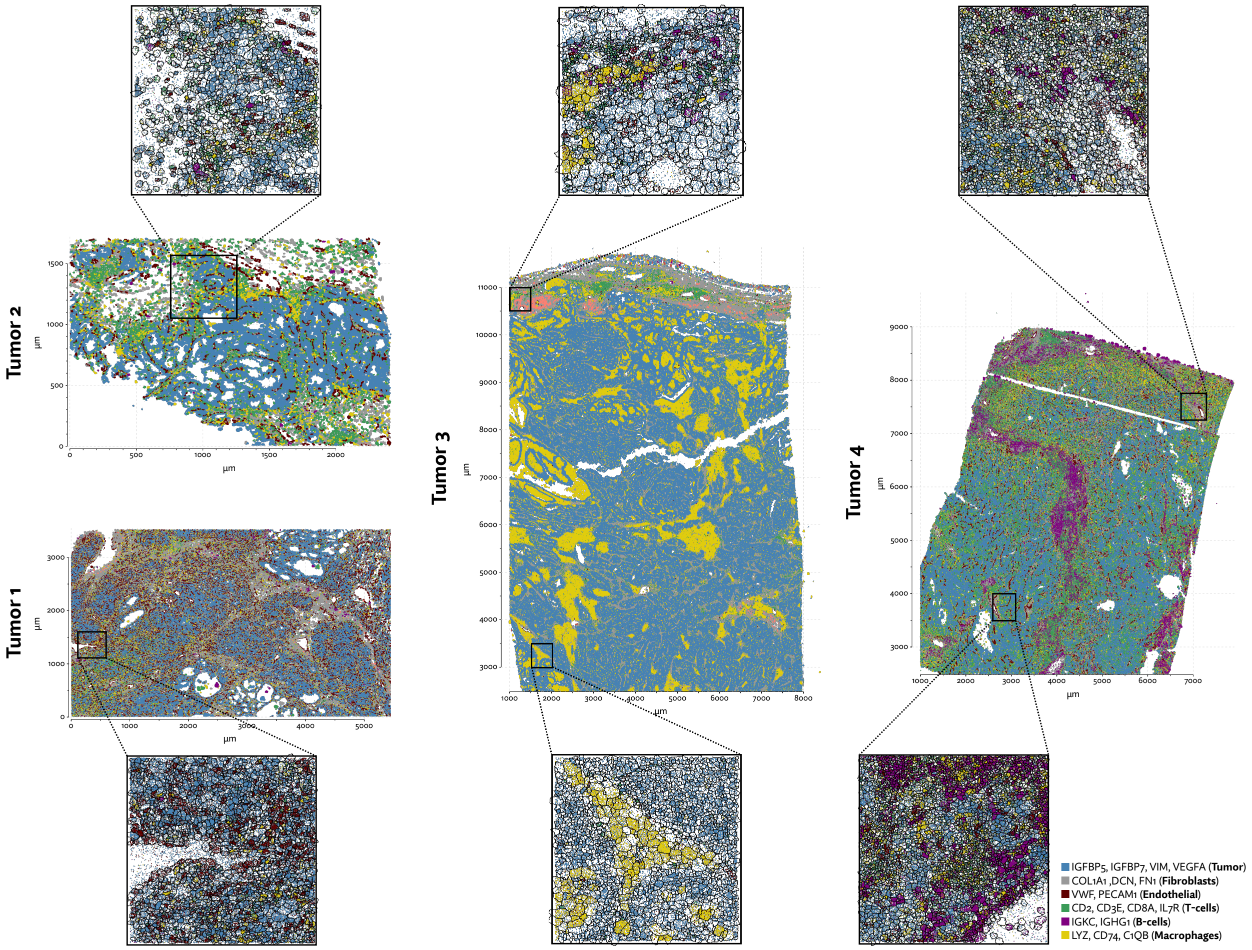


**Supplementary Figure 5:** Selected regions of the Xenium RCC dataset with annotated cell types after segmenting cells with Proseg.

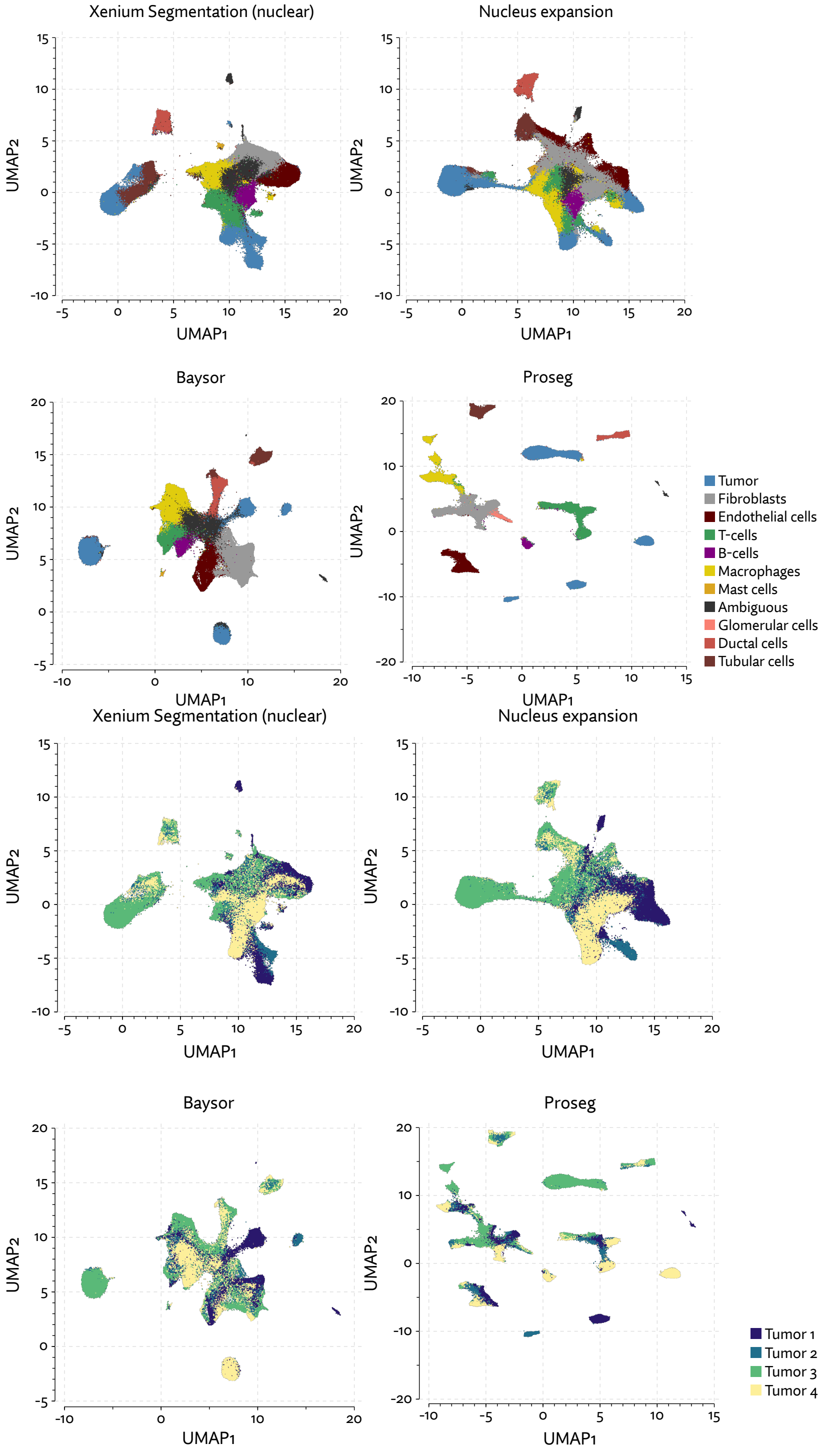


**Supplementary Figure 6:** UMAP plots for the Xenium RCC dataset using different segmentation approaches.

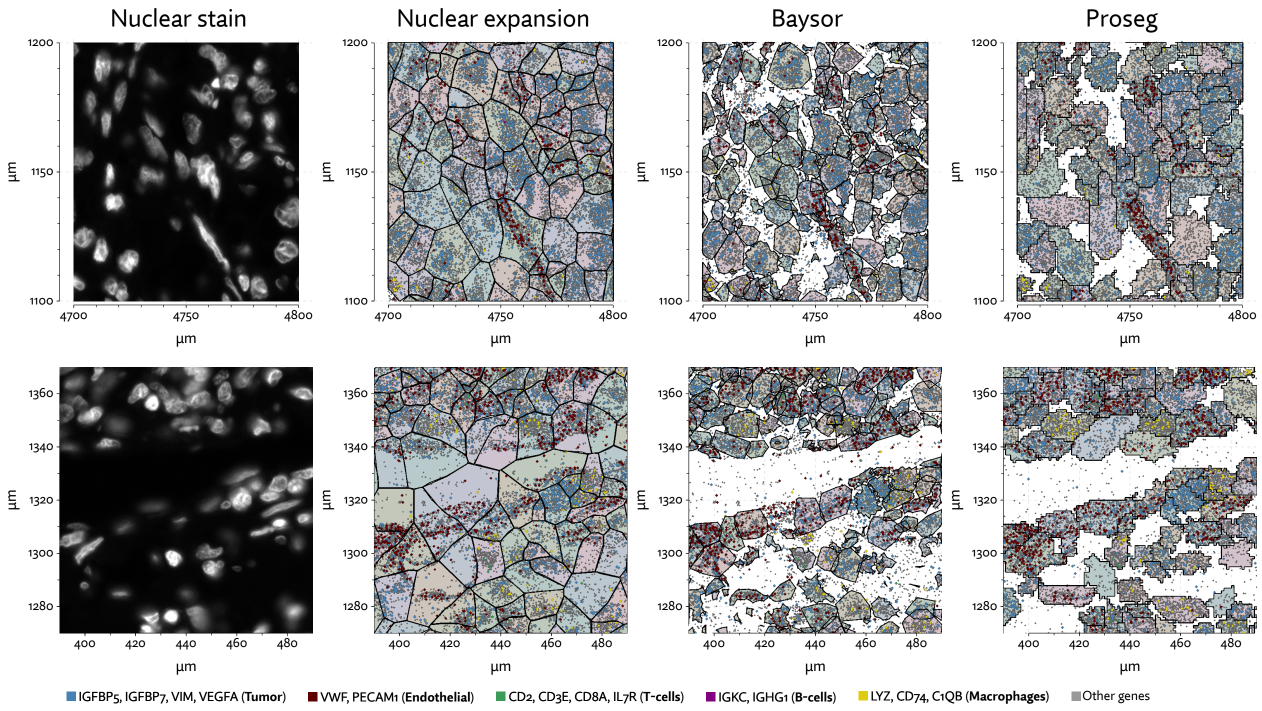


**Supplementary Figure 7:** Comparison of inferred cell boundaries in two regions of the Xenium RCC dataset, plotted alongside transcript positions and overlaid on an image of the nuclear stain.

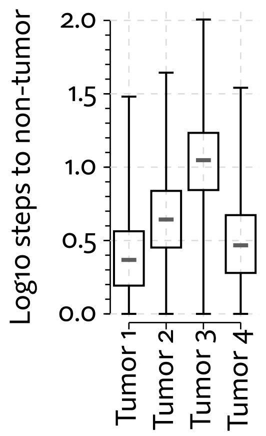


**Supplementary Figure 8:** Aggregate density of the four RCC tumor, assessed with random walk analysis.

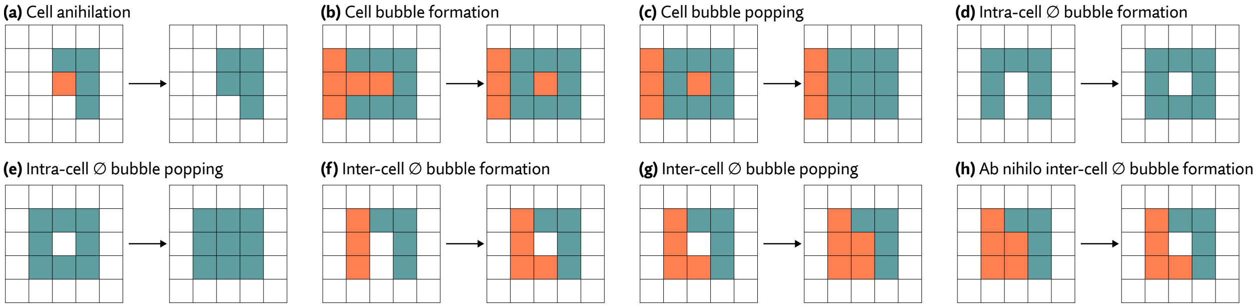


**Supplementary Figure 9:** Possible state changes that require special consideration to preserve the detailed balance property when sampling. Case (a) is irreversible, so we never propose cell annihilation. Cases (c) and (e) are irreversible, so we prohibit the formation of these bubbles in cases (b) and (d). We allow inter-cell $\varnothing$ bubble formation (f) and popping (g) by introducing an ab nihilo $\varnothing$ bubble formation proposal (h), which makes (g) reversible, preserving detailed balance.

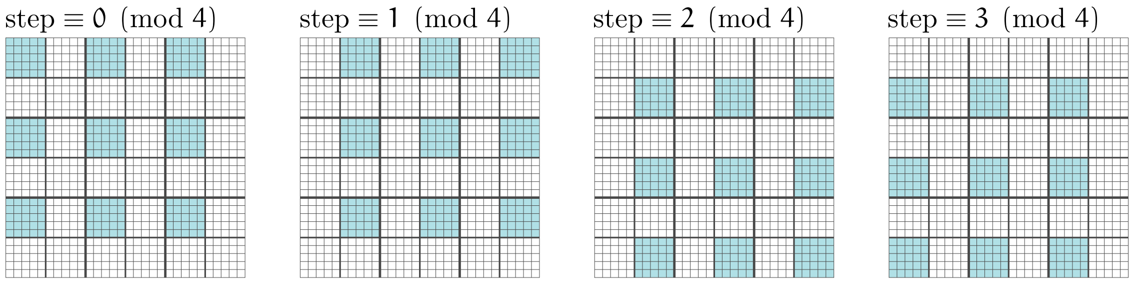


**Supplementary Figure 10:** To run the sampler in parallel on multiple CPU cores, we must avoid race conditions that can arise by simultaneously proposing multiple changes to the same cell. We handle this dividing the region into sub-quadrants and a making single proposal in each non-bordering sub-quadrant (shown in blue) on each sampler iteration.

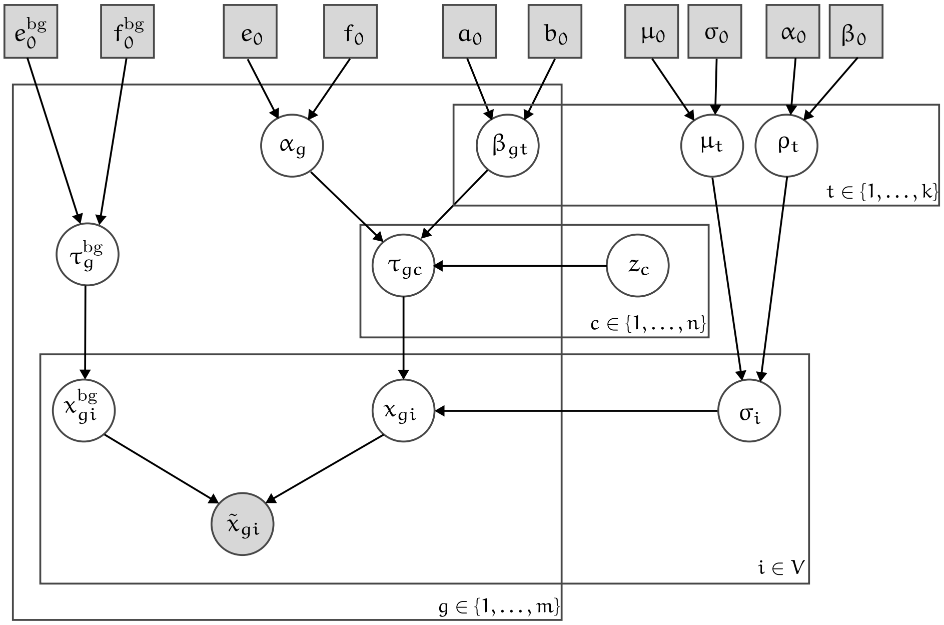


**Supplementary Figure 11:** A plate diagram of the probabilistic model of gene expression used to inform segmentation.

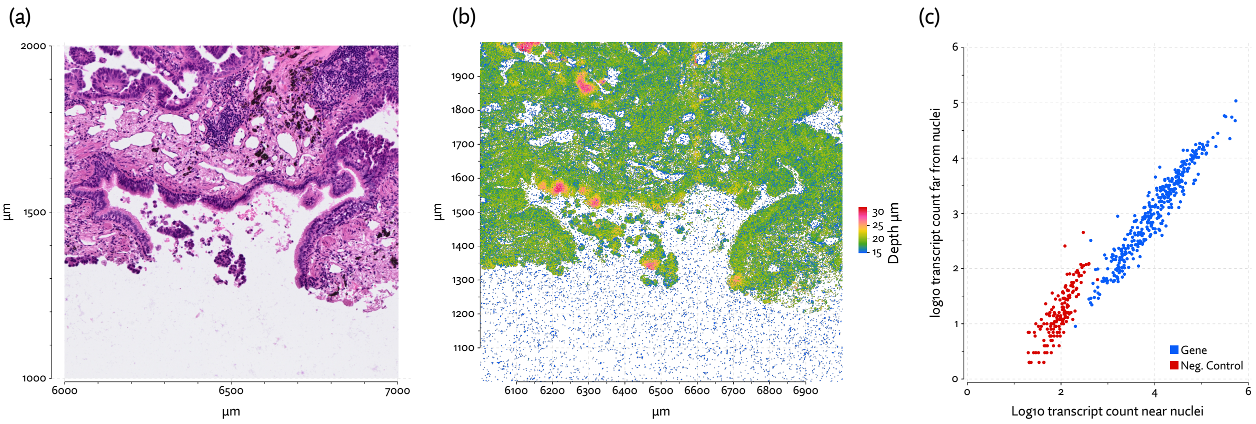


**Supplementary Figure 12:** (a) A region of the hematoxylin and eosin (H&E) stain image from the Xenium lung cancer dataset, (b) the same region showing transcripts and their measured depth (i.e. z-axis coordinate), and, (c) a comparison of the distribution of observed counts for each gene near ($\leq 50\mu m$) or and far ($> 50\mu m$) from any nuclei centroid. Negative controls, which are both barcodes with no corresponding probe and probes with no corresponding gene, are indicated in red.
